# Both Fallopian Tube and Ovarian Surface Epithelium Can Act as Cell-of-Origin for High Grade Serous Ovarian Carcinoma

**DOI:** 10.1101/481200

**Authors:** Shuang Zhang, Tao Zhang, Igor Dolgalev, Hao Ran, Douglas A. Levine, Benjamin G. Neel

**Author notes:** Corresponding authors: Benjamin G. Neel, New York University School of Medicine, 522 First Avenue, Smilow Building 12^th^ Floor, Suite 1201, New York, NY 10016.

## Abstract

The cell-of-origin of high grade serous ovarian carcinoma (HGSOC) remains controversial, with fallopian tube epithelium (FTE) and ovarian surface epithelium (OSE) each suggested as candidates. Here, by using genetically engineered mouse models and novel organoid systems, we assessed the tumor-forming capacity and properties of FTE and OSE harboring the same oncogenic abnormalities. Combined RB family inactivation (via T121 expression) and *Tp53* mutation in *Pax8+* FTE caused transformation to Serous Tubal Intraepithelial Carcinoma (STIC), which rapidly metastasized to the ovarian surface. This mouse model was recapitulated by FTE organoids, which, upon orthotopic injection, generated widely metastatic HGSOC. The same genetic lesions in *Lgr5+* OSE cells or organoids also caused metastatic HGSOC, although with longer latency and lower penetrance. Comparative transcriptome analysis was consistent with different human HGSOCs arising from FTE and OSE. Furthermore, FTE- and OSE-derived organoids showed differential sensitivity to HGSOC chemotherapeutics. Our results comport with a dualistic origin for HGSOC and suggest the cell-of-origin could influence therapeutic response.

**SIGNIFICANCE:** The cell-of-origin for high grade serous ovarian carcinoma (HGSOC) has been controversial. By generating novel GEMMs and organoid models with the same oncogenic defects, we demonstrate that HGSOC can originate from either fallopian tube epithelium (FTE) or ovarian surface epithelium (OSE). Importantly, FTE- and OSE-derived tumors differ significantly in biologic properties.

## INTRODUCTION

High-grade serous ovarian cancer (HGSOC) is the most common and deadly ovarian epithelial cancer, causing ∼70% of ovarian cancer deaths (1). Although HGSOC typically presents as a large ovarian mass accompanied by widespread peritoneal metastasis, its cell-of-origin remains controversial (2-5). Historically, as its name implies, HGSOC was thought to arise from the ovarian surface epithelium (OSE) or, more specifically, from cortical inclusion cysts, invaginations that result from normal ovulatory “wounds.” “Trapped” OSE within these cysts were believed to undergo Mullerian metaplasia and accumulate causal mutations (5-7). Cortical inclusion cysts with columnar (Mullerian) epithelia and focal p53 immunoreactivity, but not frank carcinoma, have been reported, although such lesions are relatively rare (8,9). More recently, attention has turned to the fallopian tube epithelium (FTE) as the cell of origin for HGSOC. Initial studies revealed lesions termed Serous Tubular Intra-epithelial Carcinomas (STICs) in the fallopian tube fimbria of women with *BRCA1/2* mutations undergoing risk-reducing salpingoophorectomy (10-12). Subsequently, STICs were reported in up to 60% of sporadic HGSOC patients (13-15).

Molecular analyses strongly support a fallopian tube (FT) origin of most cases of HGSOC. Essentially all HGSOCs have *TP53* somatic mutations, and STICs, primary tumors and HGSOC metastases have the same *TP53* mutation, implying a shared origin (17-20). More comprehensive genomic characterization (e.g., exome sequencing, copy number analyses, targeted amplicon deep sequencing and gene expression profiling) has shown that, in most cases, ovarian masses and distant metastases have the same “truncal” lesions as STICs, but also possess additional, often sub-clonal, genomic abnormalities, suggesting that STICs are precursor lesions (13,21). Consistent with these findings, the transcriptome of most HGSOCs more closely resembles that of normal FTE than to OSE; nevertheless, up to 12% of HGSOCs show greater transcriptional similarity to OSE (22,23). Furthermore, some genomic studies have found that HGSOC can metastasize to the FT and mimic STICs (24,25), consistent with a non-FTE origin. In addition, FTs can harbor metastases that mimic STICs but arise from other sites (e.g., uterine serous carcinoma) (25-28). The Mullerian features of HGSOC have been cited as evidence of a FT origin, as FT is a Mullerian-derived structure. However, OSE and FTE share an even earlier embryonic progenitor (6,29,30), and inappropriate expression of *HOX* family members can confer Mullerian-like features in mouse models (31).

Mouse modeling has also failed to definitively define the cell-of-origin for HGSOC. Lineage tracing identified a stem cell niche in the mouse ovarian hilum containing *Lgr5+* cells that can give rise to the entire OSE, and it was suggested that these cells were the target of transformation in HGSOC pathogenesis (32). HGSOC-like tumors have been generated by injecting adenovirus expressing Cre recombinase (Ad-Cre) into the ovarian bursae of mice bearing conditional alleles of tumor suppressor genes and/or oncogenes, including *Myc;Tp53;Brca1* (33), *Pten;Pik3ca* (34), *Tp53;Rb1* (35), or *TgK18GT*_*121*_ (N-terminal 121 amino acids of SV40 T antigen (T121) under the control of the keratin 18 promoter) and *flTp53*, ± *flBrca1 (36)*. These reports concluded that HGSOC arises from OSE, but they did not exclude the possibility that FTE cells also were infected by Ad-Cre. Conversely, a mouse model with *Pten* deletion, *Tp53* deletion or mutation, and *Brca1/2* deletion driven by the *Pax8* promoter, which is expressed selectively in FTE secretory cells (not in OSE) develops STIC-like lesions and eventually, widely metastatic HGSOC (37). Similar results were seen with in mice that express SV40 large T-antigen (TAg) (38) or with simultaneous mutation of *Brca1*, *Tp53*, *Rb1* and *Nf1* in *Ovgp1*-expressing secretory FTE (39).

In other neoplasms, the cell-of-origin can contribute to inter-tumor heterogeneity and distinct tumor biology (40-42). Therefore, we used lineage-specific Cre recombinase (Cre) lines, combined with FTE- and OSE-derived organoids, to ask whether introducing the same genetic defects into FTE or OSE results in HGSOC, and if so, whether the resultant tumors have similar properties.

## RESULTS

### Perturbing *Tp53* and the RB family in *Pax8+* cells cause STIC and metastasis

*TgK18GT121* (hereafter, “T121”) mice harbor a bacterial artificial chromosome (BAC) containing a Cre-conditional loxP-GFP-stop-loxP (LSL) T121 cassette inserted into the mouse cytokeratin (CK) 18 gene. Inducing expression of T121 (which inactivates all RB family members) by bursal injection of Ad-Cre, when combined with *Tp53* deletion or heterozygotic *Tp53*^*R172H*^ expression, results in HGSOC at a frequency of 70-90% (36). These findings were interpreted as showing that OSE gives rise to HGSOC. However, Ad-Cre injection into mice with a Cre-inducible *lacZ* allele (Rosa26-*lacZ*) resulted in mosaic LacZ staining in FTE in addition to the expected strong OSE staining (Supplementary Fig. S1), making the actual cell-of-origin in this model unclear. We therefore investigated the consequences of introducing the same ovarian cancer-associated genetic abnormalities into either FTE or OSE alone.

To this end, we generated *T121;Tp53*^*R172H*^ mice co-expressing appropriate lineage-specific *Cre* constructs. *Pax8rtTA* mice reportedly enable doxycycline (Dox)-inducible gene expression in secretory FTE (37). To confirm the lineage fidelity of this line, we generated *Pax8rtTA;TetOCre:Rosa26-tdTomato* reporter mice and *Pax8rtTA;TetOCre:Rosa26-Lacz* reporter mice (Fig. 1A, Supplementary Figure 2). Following Dox administration (0.2 mg/ml in drinking water *ad libitum*) for 1 week, histologic analysis revealed strong β-gal reactivity in the FTE, but no staining in the ovary, including the OSE (Fig. 1B). We also generated *Pax8rtTA;TetOcre;Rosa26-tdTomato* mice and exposed them to Dox for 2 days, followed by a “chase” in Dox-free water. At Day 2, Tomato+ cells co-localized with PAX8+ secretory FTE cells, as expected (Supplementary Fig. 2A). However, after 2 months of Dox-free “chase”, secretory cells (PAX8+) and ciliated cells (which stain positive for acetylated α-tubulin) both were Tomato+ (Supplementary Fig. S2B). These data confirm that PAX8+ cells give rise to ciliated cells *in vivo*, consistent with a recent study (43).

**Figure 1.**
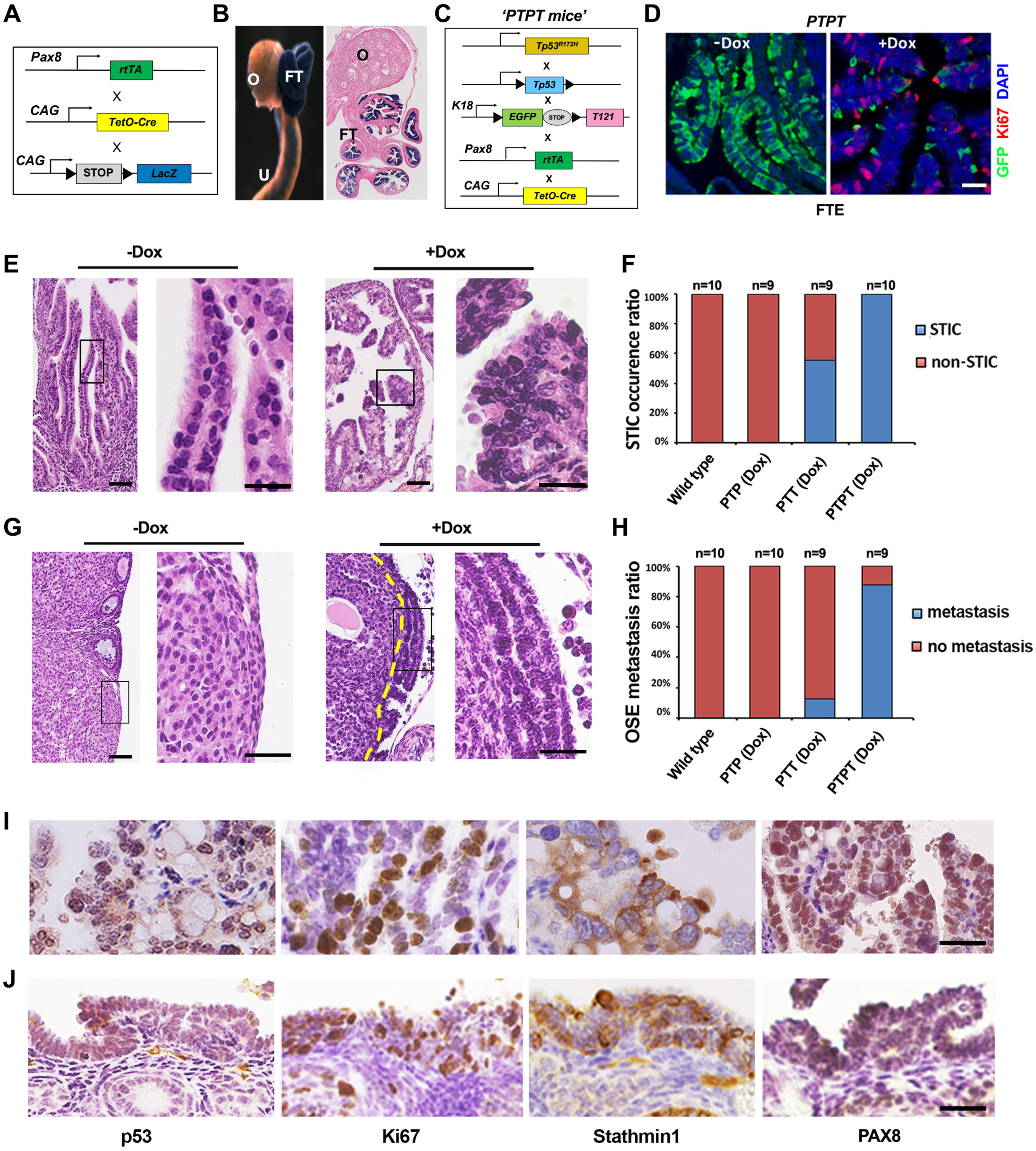
Combined *Tp53* mutation/RB family inactivation in fallopian tube epithelium results in HGSOC. **A,** Schematic showing the *Pax8rtTA*, *TetOCre*, and *Rosa26-lacZ* alleles. **B,** Whole mount (Left panel) and 10X-magnified histological section (right panel) of genital tracts of *Pax8rtTA*;*TetOCre*;*Rosa26-LacZ* female mice, treated with Dox (2 mg/ml in drinking water for 2 weeks), stained with X-gal to detect β-galactosidase and counterstained with eosin. **C,** Schematic of *Pax8rtTA;TetOcre; Tp53*^*R172H/-*^*;T121* (PTPT) mice. **D,** Immunofluorescence staining for GFP (marker for keratin 18+ FTE cells, green), Ki67 (proliferation marker, red) and DAPI (nuclear stain, blue) with or without Dox trea™ent as in **B**. Scale bar: 10 µm**. E,** Representative H&E-stained section of fallopian tube from PTPT mice with or without Dox (2mg/ml in drinking water for 1 month). Boxed regions in left panels are magnified in the right panel of each pair. Scale bar: 50 µm. **F,** % mice of each genotype with STIC, assessed after 1 month of Dox trea™ent. **G,** Representative H&E-stained ovary sections from PTPT mice, without or with Dox trea™ent for 1 month. Boxed regions in left panels are magnified in the right panel of each pair. The yellow dashed line shows the border between OSE and underlying cells. Arrows show areas of neoplasia on ovarian surface. Scale bar: 50 µm. **H,** % of mice of each genotype that developed ovarian metastasis, assessed after 1 month of Dox trea™ent. **I** and **J**, IHC for key STIC markers in representative sections of FTE (top) and OSE (bottom) of PTPL mice. Scale bar: 10 µm.

Having confirmed the selective expression of *Pax8rtTA* in FTE, we generated *Pax8rtTA;TetOcre;Tp53*^*R172H/fl*^ (hereafter, “PTP mice”), *Pax8rtTA;TetOcre;T121* mice (hereafter, “PTT mice”), which direct inducible expression of T121 alone in *Pax8+* cells, and *Pax8rtTA;TetOcre;Tp53*^*R172H/fl*^*;T121* mice (hereafter, “PTPT mice”), which enable Dox-dependent expression of T121 and deletion of one *Tp53* allele on a background of *Tp53*^*R172H*^ heterozygosity. As noted above, T121 mice contain a *GFP* reporter embedded within their LSL cassette (Fig. 1C), which allows deletion efficiency to be monitored by absence of GFP. As expected, in the absence of Dox, GFP was detected diffusely in FTE, which exhibited very low levels of proliferation, as shown by Ki67 staining. Following Dox administration, the newly generated GFP-cells showed substantial Ki67 positivity, indicating that T121 evokes FTE proliferation (Fig. 1D). Within 1 month, FTE in Dox-activated PTPT mice showed clear evidence of transformation, with characteristic features of STIC, including secretory cell proliferation, loss of polarity, cellular atypia, and serous histology (Fig. 1E and F). As early as 1 month post-Dox, metastases were detected on the ovarian surface (Fig. 1G, compare boxed regions). Notably, 5/9 PTT mice underwent similar FTE transformation, suggesting that RB family inactivation alone can promote STIC. This finding is consistent with a previous study in which expressing SV40 large T antigen under control of the *Ovgp-1* promoter resulted in STIC (38). We did not detect STIC in PTP mice (0/9), but combining *Tp53* perturbation with T121 expression significantly increased FTE transformation (from 55% to 100%) and metastasis (from 0 to 88%) to the ovary (Fig. 1F and H). Immunohistochemical analysis (IHC) showed HGSOC-related features, including TP53 accumulation, proliferation and PAX8 expression in the primary lesion and the ovarian metastases of PTPT mice (Fig. 1I and J).

Unexpectedly, PTT and PTPT mice died within 2 months (Supplementary Fig. S3A), but their demise was not due to peritoneal masses/obstruction, as would be expected with lethal HGSOC. Instead, these mice had markedly enlarged thymi, which led to breathing difficulties and ultimately, respiratory arrest (Supplementary Fig. S3B). Combined *Rb/p130* deletion/*p107* heterozygosity causes a similar phenotype (44), and surprisingly, re-analysis of *Pax8rTtA;TetO-Cre; Rosa 26-tdTomato* mice showed that *TetO-Cre* is active in thymic epithelial cells (although not in FTE) even in the absence of Dox (Supplementary Fig. S3C). This unanticipated “leakiness” of *TetO-Cre* in the thymic epithelium precluded us from asking if *Pax8-*driven T121 expression/*Tp53* perturbation leads to widespread metastasis, although the early ovarian studding and our organoid studies (see below) make this highly likely.

### FTE organoids recapitulate fallopian tube differentiation and transformation *in vitro*

Organoids initiated from adult stem cells can often be propagated long-term and are useful for cancer modeling (45,46). Culture conditions for human FTE organoids have been established (47), but OSE organoids have not been reported, nor have human or mouse organoids been used to model HGSOC. We developed serum-free defined media that allow mouse FTE and OSE (see below) organoids to be passaged indefinitely (>30 passages) in Matrigel-based media, cryopreserved, and re-established in culture (see Methods). By 48hr after seeding FACS-purified EPCAM-/CD45-cells into such media, small (1-5 µm in diameter), round, cystic spheres appeared (EPCAM+CD45-), and they expanded rapidly to form large (100-1000 µm in diameter) hollow cysts. After 10 days, epithelial invagination resulted in mucosal folds, morphologically recapitulating FT architecture (Fig. 2A). Organoids cultured longer than 7 days contained secretory and ciliated cells, like their tissue of origin (Fig. 2B).

**Figure 2.**
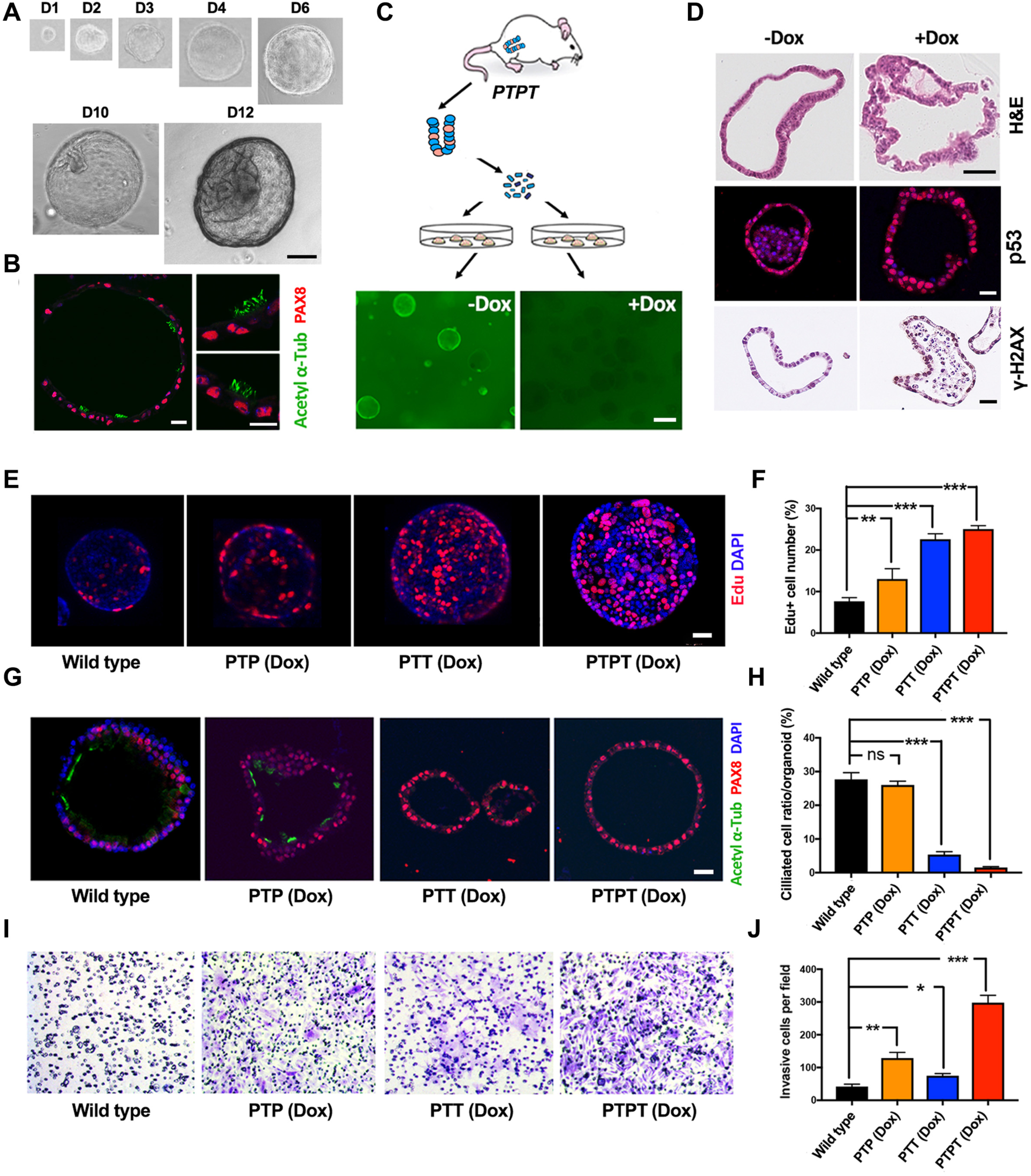
Establishment and characterization of FTE organoids. **A,** Bright field image of organoid developing from single FTE cell at the indicated days of culture. Scale bar, 20 µm. **B,** Immunofluorescence staining for ciliated cell marker acetyl-α-tubulin (green) and secretory cell marker PAX8 (red) in FTE organoid after 7 days in culture. Scale bar, 20 µm. **C,** Top: Schematic showing generation of FTE organoids from PTPT mice. Bottom: GFP in organoids with or without Dox trea™ent; Dox (500 ng/ml) was added after seeding cells and GFP was gone in PTPT organoids after 2 passages. Scale bar, 10 µm. **D,** Sections of PTPT organoids after 2 consecutive passages, with or without Dox trea™ent (500 ng/ml), stained with H&E or subjected to immunofluorescence or IHC staining for the indicated markers. Scale bar, 20 µm. **E**, Organoids were established from the indicated mice, incubated with EdU (2 µM) and DAPI (1µg/ml) for 2 hours on day 7 of culture, and visualized by immunofluorescence. Scale bar, 20 µm. **F,** % EdU-positive cells in organoids established from the indicated mice. **G**, Representative immunofluorescence images of PAX8 (red) and acetylated-α-tubulin (green) staining of the indicated organoid lines at day 7 of culture. Scale bar, 20 µm. **H,** % ciliated cells (aceylated-α-tubulin+) in organoids established from the indicated mice. **I,** Representative images of the bottom surface of Transwell units seeded with the indicated organoids. **J,** Quantification of invasive cells in **I**. Data represent means ± SEM from three mice of each genetic background. *P<0.05, **P<0.01,***P<0.001.

To evaluate the utility of this system for modeling HGSOC, we established organoid cultures from FTE of wild type, PTP, PTT, and PTPT mice. Dox-treated PTT and PTPT organoids activated *Cre* expression, as indicated by loss of GFP after 3 passages (Fig. 2C and data not shown). Dox-treated (but not control) PTPT organoids exhibited frank dysplasia, TP53 accumulation, and staining for γH2AX, a marker of DNA damage (Fig. 2D). Consistent with our GEMM results (Fig. 1), PTT and PTPT organoids proliferated more rapidly and were larger than controls (Fig. 2E and F, Supplementary Fig. S4A and B). By contrast, organoid-forming efficiency was not increased in mutants, suggesting that *Tp53* mutation and/or RB family inactivation did not affect FTE self-renewal, as least under our culture conditions (Supplementary Fig. S4C). STIC and HGSOC typically are PAX8+ (although staining can be mosaic), and they do not express ciliated cell markers (48). Similarly, ciliated cells were undetectable in PTT or PTPT organoids, suggesting that the RB family inactivation restricts FTE differentiation to the secretory lineage (Fig. 2G and H). *Tp53*^*R172H/-*^ mutation did not impair differentiation, but it enhanced invasive capability (Fig. 2I and J). Thus, FTE organoids can be used to deconstruct various aspects of the malignant phenotype and attribute each to specific genetic defects.

### FTE-derived organoids can give rise to HGSOC

We previously found that primary human HGSOC cells implanted into the mouse mammary fat pad (MFP) recapitulate HGSOC histomorphology, inter- and intra-tumor heterogeneity, and patient drug response (49,50). Therefore, as an initial test of their tumor-forming capacity, we injected wild type and mutant organoids into the MFPs of *nu/nu* mice (Fig. 3A). Wild type and *Tp53*^*R172H/-*^ organoids formed glandular structures composed of simple cuboidal cells that persisted for at least 4 weeks, but these disappeared (n=0/8 for wild type, n=0/9 for PTP organoids) by 2 months (Fig. 3B-D). By contrast, PTPT organoids formed easily palpable (>2mm) masses of high grade, poorly differentiated adenocarcinoma with prominent cellular/nuclear pleomorphism and frequent mitoses (Fig. 3D). PTT organoids also gave rise to HGSOC in 3 out of 8 mice, but consistent with our finding that biallelic *Tp53* perturbation is associated with disease progression (Fig. 1F and H), PTPT organoids were more tumorigenic (7 out of 8 mice, Fig. 3B). The ovary is thought to provide trophic signals that enable HGSOC growth and metastasis (51), so we also assessed tumorigenicity of organoids injected into the ovarian fat pad. PTT and PTPT organoids both gave rise to primary tumors, but within 3 months, 5/7 mice injected with PTPT organoids developed ascites, sometimes hemorrhagic, with widespread peritoneal studding similar to the human disease. By contrast, 0/7 lines of PTT-derived organoids showed evident metastasis at 9 months post-injection (Fig. 3E and F). Metastases expressed HGSOC markers, including PAX8, p53, Ki67, γH2AX, Stathmin1 and p16 (Fig. 3G). Hence, our FTE organoid system recapitulates HGSOC biology morphologically and molecularly.

**Figure 3.**
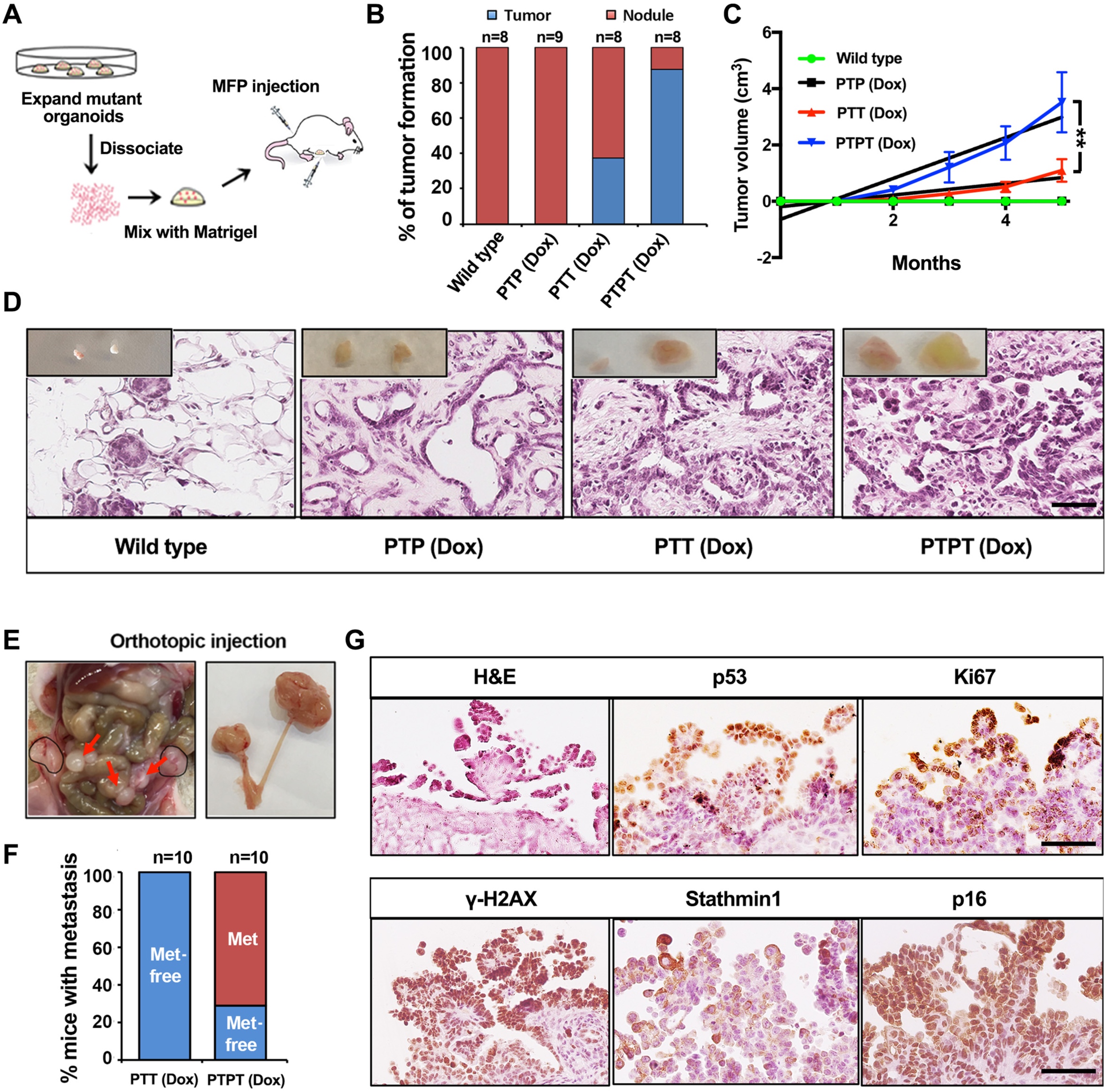
Transplanted organoids recapitulate HGSOC progression and metastasis. **A,** Scheme depicting organoid transplantation. **B,** % of tumors formed in mammary fat pads within 6 months after injection of 10^5^ cells from wild type, PTP (Dox-treated), PTT (Dox-treated), and RPTP (Dox-treated) organoids, respectively. **C,** Graph showing the average tumor volume of mice injected with organoids of the genotypes indicated in **B**. Data represent mean ± SE, **P<0.01. **D,** H&E-stained sections of mammary fat pads of mice injected with 10^6^ cells from the indicated organoids at 3 months after transplant. Inserts show gross appearance of nodules/tumors. Scale bar, 50 µm. **E,** Left panel: exposed mouse abdomen 3 months after orthotopic injection of mice with 10^6^ PTPT organoid cells, showing large ovarian mass (circled on left panel) and widespread peritoneal metastasis (arrows). Right panel: the genital duct dissected from the left panel shows large tumors on ovary. **F,** % of mice injected with the PTT (dox) organoids (0%, n=10) and PTPT (Dox) organoids (30%, n=10) that develop peritoneal metastasis, assessed at 6 months following injection of 10^6^ organoid cells. **G,** H&E-stained sections and IHC staining for the indicated HGSOC markers in omental metastases. Scale bar, 50 µm.

### OSE also can give rise to HGSOC

Ad-Cre injection into the ovarian bursa of *Tp53*^*R172H/fl*^*;T121* mice results in HGSOC (36), but as shown above, FTE cells also are infected under these conditions. As an initial test of whether OSE itself could serve as the HGSOC cell-of-origin, we performed salpingectomies 3 days after Ad-Cre injection into *Tp53*^*R172H/fl*^*;T121* mice (Supplementary Fig. S5A). Gross inspection and histology of the ovaries of injected mice confirmed that the FT had been removed (Supplementary Fig. S5B). Nevertheless, when mice were examined 3 months later, the OSE showed abnormal morphology and hyperproliferation (Supplementary Fig. S5C), which prompted us to more carefully evaluate OSE as a potential cell-of-origin for HGSOC.

*Lgr5+* embryonic and neonatal populations establish the OSE lineage and the fimbrial FTE (52). In adult mice, however, *Lgr5* expression is concentrated in the ovarian hilum, together with the stem cell markers CD133 and ALDH, and lineage-tracing using *Lgr5Cre*^*ERT2*^ mice showed that *Lgr5+* cells can repopulate the entire OSE (32). To confirm the lineage fidelity of this *Cre* line, and in particular, to assess its expression in adult FTE, we generated *Lgr5Cre*^*ERT2*^*;Rosa26-tdTomato* reporter mice. On Day 2 after administration of a single dose of 4-OHT (to induce CRE activity), there were sparse Tomato+ cells in OSE. By 4 months, though, these cells had expanded and populated a significant fraction of OSE. By contrast, there was no FTE expression at either time (Fig. 4A and B).

**Figure 4.**
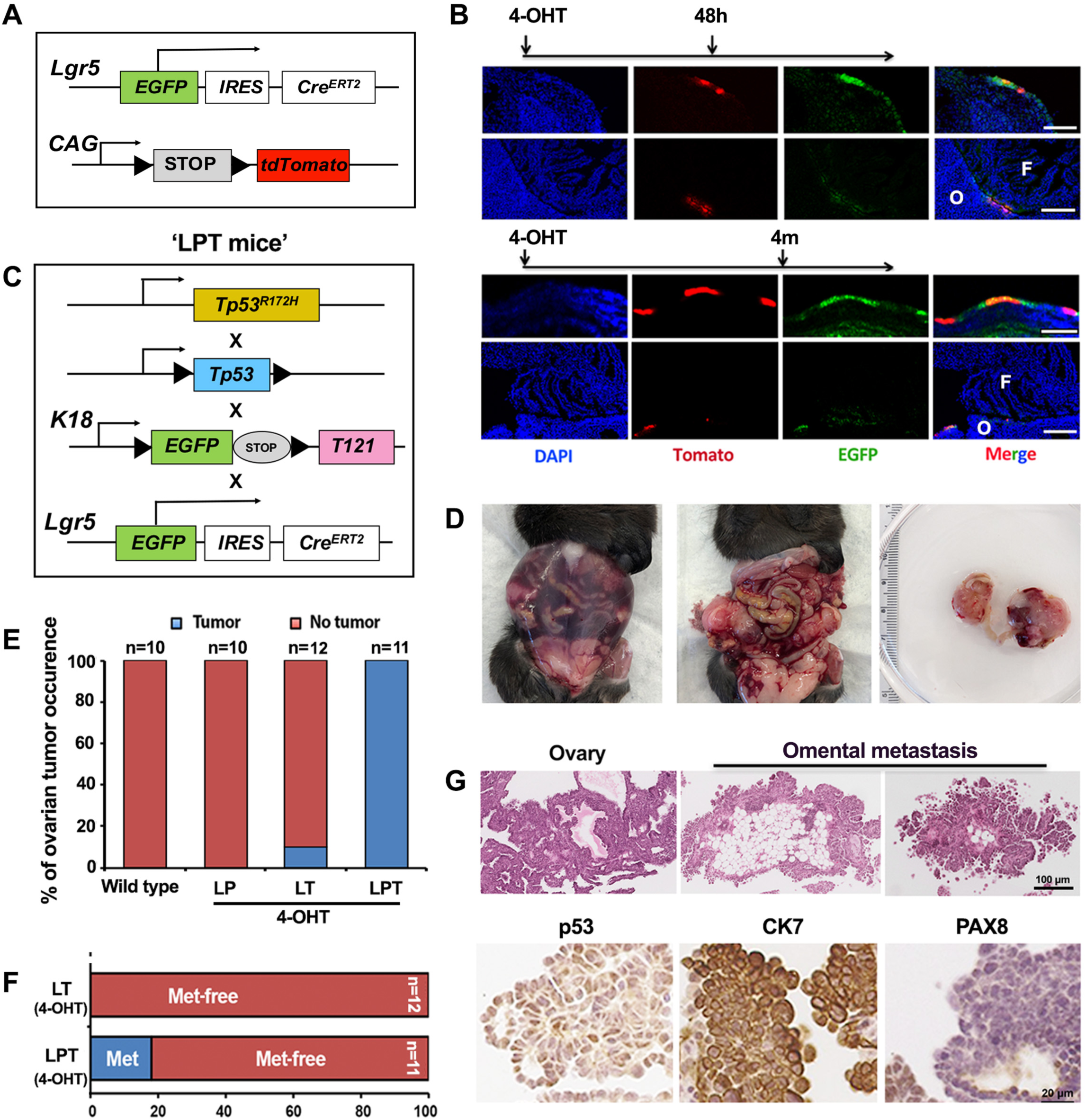
Combined *Tp53* mutation/RB family inactivation in *Lgr5+* OSE also causes HGSOC-like malignancy. **A,** Scheme depicting *Lgr5Cre*;*Rosa26-tdTomato* mice.**B,** EGFP co-immunostaining of Tomato+ OSE clone in ovary and fallopian tube sections, 48 h or 4 months post 4-OHT induction of *Lgr5-Cre;Rosa26-tdTomato* mice. Scale bars, 50 µm. **C,** Scheme depicting generation of *Lgr5Cre*^*ERT2*^;*T121*;*Tp53*^*R172H/-*^ mice (LPT mice). **D,** Exposed abdominal cavity of an LPT mouse 11 months after 4-OHT trea™ent, showing marked abdominal distention due to ascites, large ovarian tumors and peritoneal studding. **E,** % of mice of the indicated genotypes showing ovarian tumors at 18 months post 4-OHT induction. **F,** % LT and LPT mice with peritoneal metastasis 18 months after 4-OHT trea™ent. **G,** Representative p53, CK7 and PAX8 staining (by IHC) in metastatic tumor from an LPT mice.

We next generated *Lgr5-Cre*; *Tp53*^*R172H/fl*^*;T121* (LPT) mice, and compared them with *Lgr5-Cre*; *Tp53*^*R172H/fl*^ (LP) and *Lgr5-Cre*;*T121* (LT) mice, respectively (Fig. 4C). All mice received a single intraperitoneal dose of 4-OHT. As expected, LP mice (n=0/10) did not develop tumors, and only 1 out of 12 LT mice showed an obvious ovarian mass by 18 months after 4-OHT trea™ent. By contrast, all (11/11) LPT mice had large, palpable, abdominal masses by 11 months post-4-OHT injection (Fig. 4D and E). Gross inspection revealed markedly enlarged, hemorrhagic ovaries, with 16% (n=2/11) developing multifocal peritoneal carcinomatosis and ascites (Fig. 4D and F). Histology and IHC revealed poorly differentiated adenocarcinoma with papillary and micropapillary/filigree morphology (Fig. 4G), resembling human HGSOC. Notably, however, these tumors showed little PAX8 staining.

As an additional test of whether *Lgr5+* OSE cells directly evoke HGSOC transformation, we generated compound LPT;*Rosa 26-tdTomato* mice (Supplementary Fig. 6A), administered a single dose of 4-OHT, and “chased” for 3, 6, or 9 months (Supplementary Fig. 6B). By 3 months following CRE activation, neoplasia had developed from *Lgr5+* cells (Supplementary Fig. 6C). These lesions expanded, and at 6 months, >50% of the ovarian surface was covered by malignant, Tomato+ cells, whereas at 9 months, the surface was almost entirely overrun (Supplementary Fig. 6C-F). Histologic examination again showed multi-villus neoplasia (Supplementary Fig. 6D and G), and IHC revealed highly proliferative tumors expressing HGSOC markers, including Wilms’ tumor 1 (WT1) and Stathmin1 (Supplementary Fig. 6G). Neither hyperproliferation nor neoplasia was evident in the FT of LPT mice (Supplementary Fig. S7A), and the OSE-derived tumors also were PAX8-negative (Supplementary Fig. S7B).

### OSE-derived organoids also give rise to ovarian cancer

We were unable to culture OSE organoids in the FTE medium. However, with additional factors, such as hydrocortisone, estrogen, LIF and forskolin, organoids could be propagated from single OSE cells (Fig. 5A). These organoids expressed epithelial markers, such as E-cadherin, but unlike FTE organoids, they did not express PAX8 (Fig. 5B). As noted above, *Lgr5+* cells can repopulate the entire OSE, suggesting that they have stem/progenitor activity. Consistent with this notion, GFP^hi^ (*Lgr5*-expressing) cells accounted for almost all organoid-forming capability, compared with GFP^neg^ (*Lgr5-negative*) cells (P<0.001, Fig. 5C). We next generated OSE organoids from wild type, LP, LT and LPT mice, and added 4-OHT to the medium for 48h to activate CRE. Following 4-OHT trea™ent, T121 was expressed in LT and LPT organoids, as expected (Fig. 5D). Organoid size was not affected, but mutant organoids were more condensed, containing larger numbers of cells (Fig. 5E). Although organoid-forming capacity was similar among the 4 groups, LPT (P<0.001), and to a much lower extent (P<0.01), LT organoids grew faster (Fig. 5G). Finally, we implanted all four types of organoids into the MFP and orthotopically (Fig. 5H). Consistent with the cognate GEMMs (Fig.4F), only LPT organoids induced metastasize tumors, and organoids and the tumors they evoked had similar histology and marker expression (Fig. 5I and J).

**Figure 5.**
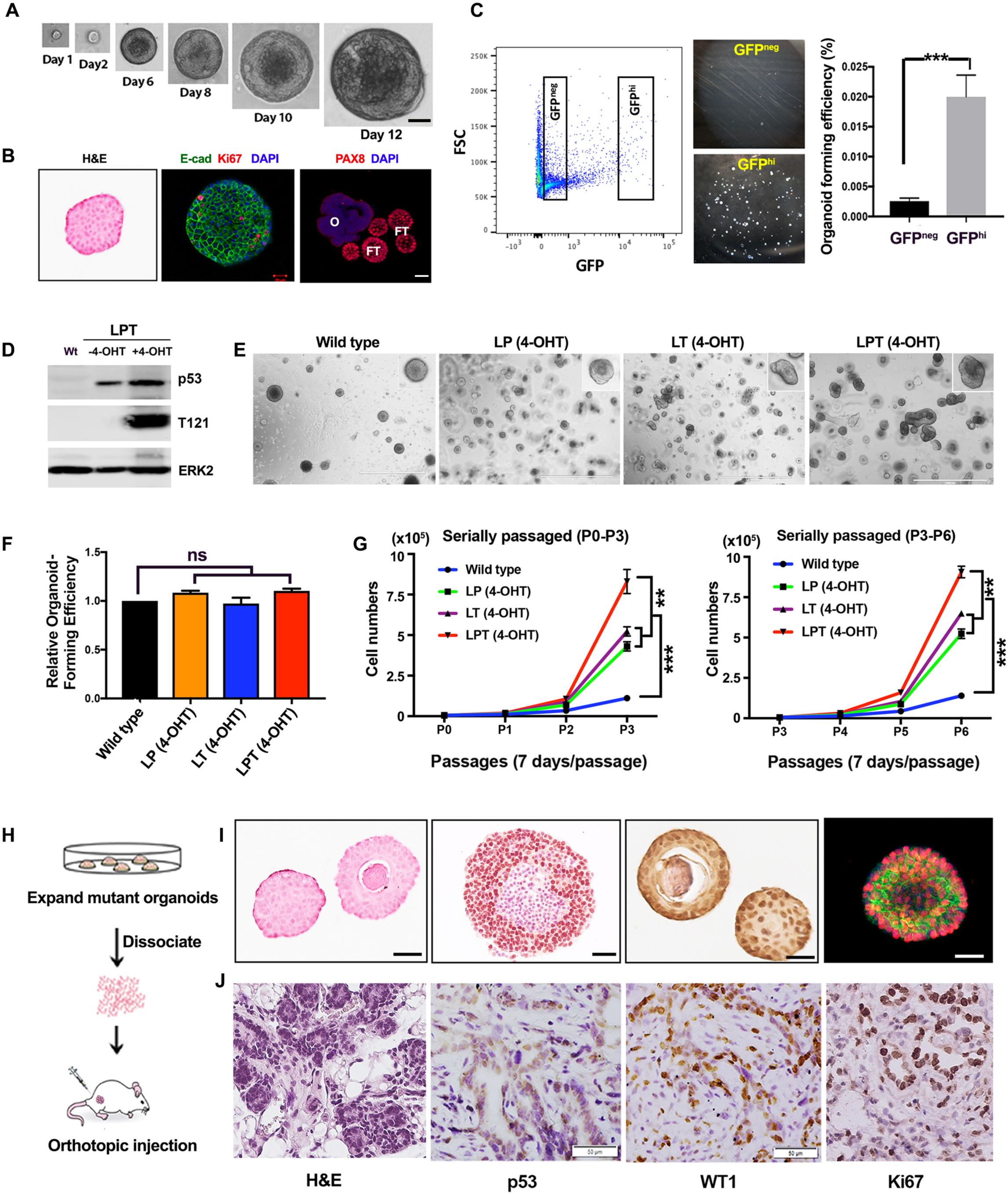
Transplanted OSE organoids also can give rise to HGSOC. **A,** Representative serial images of an OSE organoid at the indicated times after seeding. Magnification: ×10. Scale bar, 20 µm. **B,** Left panel, H&E stain of an OSE organoid. Middle panel, Immunofluorescence staining for E-cadherin (E-cad, green) and Ki67 (red). Right panel, immunofluorescence staining for PAX8 in a mixed culture of FTE (FT) and OSE (O) organoids. Note that only FTE organoids are strongly PAX8+. Scale bar, 20 µm. **C,** Representative flow cytometric plot, showing gates used to purify EGFP^hi^ and EGFP^neg^ populations from *Lgr5-GFP* OSE cells (Left panel), and bright field pictures, showing organoids that developed from FACS-purified cells (5,000 cells/well) cultured for 6 days (Middle panel). Right panel shows organoid-forming efficiency of the two populations. (n?=?10 wells per group combined from three experiments). Data indicate means ± SEM, ***p < 0.001. **D,** Representative immunoblot for T121 in LPT organoids without 4-OHT trea™ent and 2 weeks after 4-OHT induction. ERK2 serves as a loading control. WT, wild type organoid. **E,** Micrographs showing typical OSE organoids from mice of the indicated genotypes after 6 days in culture. **F,** Relative organoid-forming efficiency of OSE cells from mice of the indicated genotypes. Cells (5000) were seeded from the 3^rd^ passage of each genotype, and organoids were counted at day 6 of culture. For each genotype, 3 independent mice were used to generate organoids. Data represent means ± SEM. Statistical analysis was performed by Dunnett’s multiple comparison test. ns, not significant. **G,** Growth curves of OSE organoids from the indicated mice. Note that cultures show exponential growth within the time window analyzed. Cells per well were counted at each passage from P1-P3 and P4-P6. Organoids from 3 independent mice for each group were used, each in duplicate. Data represent means ± SEM. Statistical analysis was performed by Dunnett’s multiple comparison test. **P<0.01, ***p < 0.001. **H,** Scheme depicting OSE organoid transplantation (**I** and **J**) Representative H&E staining and IHC for the indicated markers in LPT organoids (top) and the orthotopic tumors derived from these organoids, 4-months post injection of 10^6^ cells (bottom).

### Transcriptome analysis reveals similarities and differences between FTE- and OSE-derived HGSOC

To molecularly characterize our HGSOC mouse models in an unbiased manner, we performed RNAseq on FTE-derived tumors (T-FT hereafter) from PTPT mice and OSE-derived tumors (T-O hereafter) from LPT mice. Pooled normal OSE (N-O hereafter) and FTE (N-FT hereafter) samples (3 each) were profiled as controls. Unsupervised analysis showed that FTE-derived (tumor and normal) samples segregated from OSE-derived (tumor and normal) samples (Fig. 6A). Multiple genes were differentially expressed in the two normal cell types (N-O vs N-FT) and in the tumors that derive from them (T-O vs T-FT); indeed, nearly 3,000 lineage-specific genes contributed to significant differences in gene expression (FDR-adjusted P ≤ 0.05) between FTE-and OSE-derived tumors, respectively (intersection between the N-O vs. N-FT and T-O vs. T-FT ovals in the Venn diagram in Fig. 6B; also see Supplementary Table S1A). Furthermore, supervised analysis revealed that the top 50 and top 100 differentially expressed genes between FTE-derived and OSE-derived tumors largely reflected differential gene expression in N-FT and N-O, although other (non-lineage) genes contribute to the top 1,000 DEGs (Supplementary Fig. S8). Comparing N-FTE and N-O, *Pax8* (logFC=4.05, P_adj_=5.5×10^−14^), *Ltf* (logFC=-5.92, P_adj_<1×10^−15^), *Slc34a2* (logFC=4.84, P_adj_<1×10^−15^), which are known FT-specific genes (53), are highly expressed both in N-FT and T-FT, compared with N-O and T-O. By contrast, the known OSE-specific gene *Lgr5* (logFC=5.54, P_adj_=0) was expressed at significantly higher levels in N-O and T-O than N-FT and T-FT, as were *Nr5a*1 (encoding a transcriptional activator involved in sex determination and differentiation of steroidogenic tissues, logFC=6.5, P_adj_<1×10^−15^), *Gata4* (encoding a zinc finger TF), *Lhx9* (encoding a target of GATA4, logFC=6.58, P_adj_<1×10^−15^), and *Unc45b* (encoding a GATA4 chaperone, logFC=6.6, P_adj_<1×10^−15^); see Supplementary Fig. S8 and Supplementary Table 1B).

**Figure 6.**
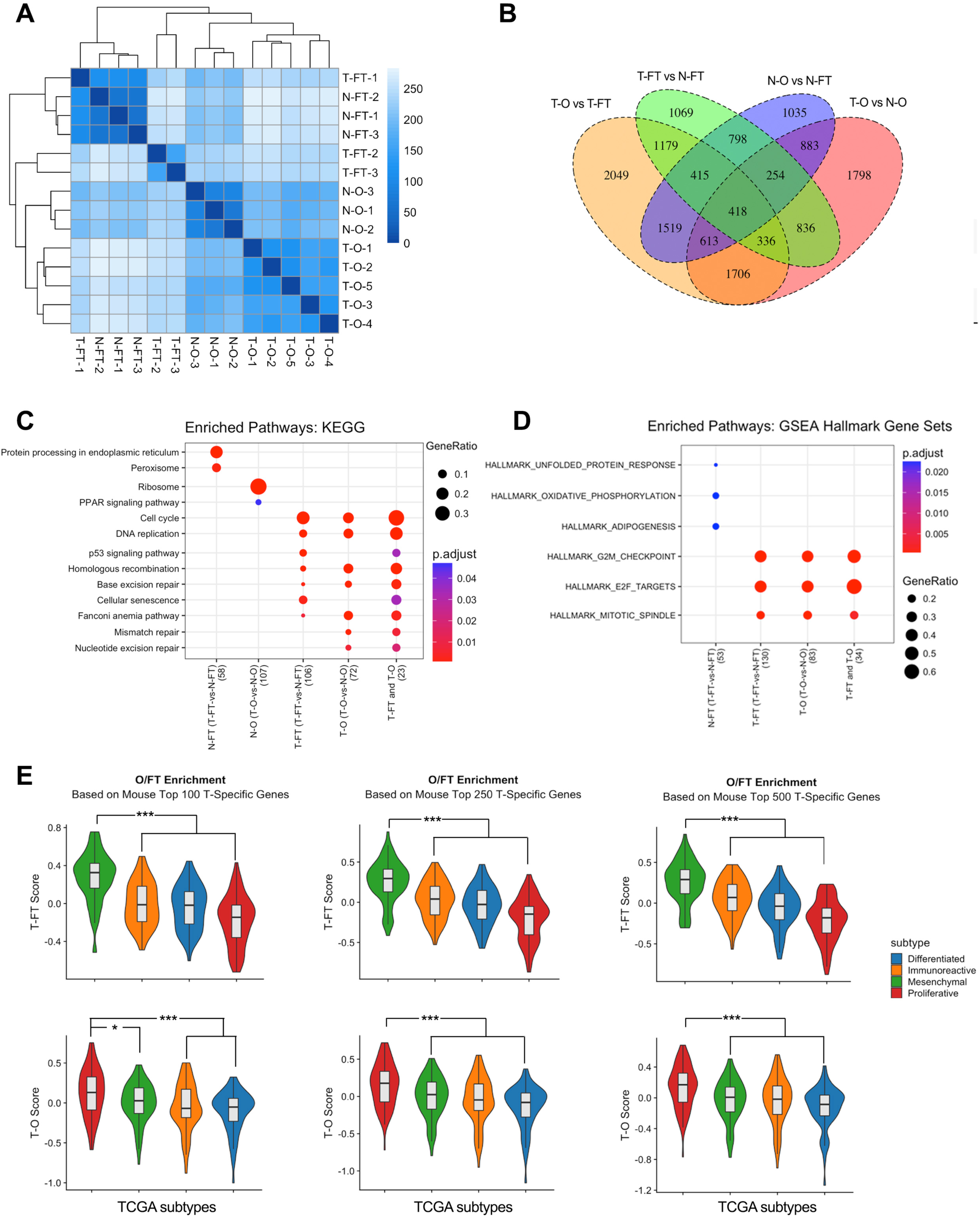
Comparative transcriptome analysis of mouse and human HGSOC. **A,** Hea™ap of sample distances with hierarchical clustering based on the overall gene expression levels of normal FTE (N-FT), normal OSE (N-O), and tumors derived from FTE (T-FT) and OSE (T-O), respectively. The cell shading represents the Euclidian distance for each sample pair. B, Venn diagram showing number of differentially expressed genes (adjusted P <0.05) in T-FT vs N-FT, N-O vs N-FT, T-O vs N-O, and T-O vs T-FT. **C and D,** Significant KEGG pathway (C) and GSEA (**D**) analyses of the top 250 differentially expressed genes (DEGs) between the indicated groups **E,** Application of Ovarian or Fallopian tube (O/FT) enrichment score, based on mouse Top 100, 250 and 500 DEGs (T-O or T-FT, respectively), to TCGA samples. Color coding indicates the transcriptional subtype assigned to each sample by TCGA. See Methods for details. T-FT and T-O signatures were compared across HGSOC molecular subgroups. The T-FT score was highest in the mesenchymal subgroup (upper panel); the T-O signature was highest in the proliferative subgroup (lower panel). Statistical analysis was performed by Dunnett’s multiple comparison test. *p<0.05, ***p < 0.001.

Next we searched for common DEGs in each sample type. Nearly 7,000 (6844) DEGs were seen in OSE-derived tumors compared with normal OSE, whereas 5305 DEGs were yielded in T-FT vs N-FT (all at the P<0.05 level). Of these, 1844 genes were shared between T-FT and T-O, compared with the cognate normal samples (Supplementary Fig. S8 and Supplementary Table S1C). KEGG analysis showed that DEGs in normal FTE, compared with normal OSE, were enriched for different pathways; e.g., peroxisome-related transcripts were enriched in the FTE, whereas ribosome-related genes, were enriched in the OSE. Although a substantial fraction of differential gene expression in T-FT vs T-O reflected lineage-specific gene expressions, tumors also shared multiple DEGs (compared with their cognate normal cells-of-origin). These were enriched for genes involved in pathways annotated as controlling the cell cycle, DNA replication and DNA repair (Fig.6C). Of note, the “p53 signaling pathway” was more enriched in the FTE-derived tumors, whereas homologous recombination, mismatch repair and nucleotide excision repair were more enriched in OSE-derived tumors (Fig. 6C). Similar to the KEGG analysis, the most enriched GO categories in all of the tumor sets involved cell/nuclear division and cell cycle control and DNA replication and repair, while none of the pathways enriched in the normal FT and normal OSE (Supplementary Fig. S9A). GSEA showed a predominance of G2/M checkpoint control and E2F target genes in all of the tumor samples (Fig. 6D and Supplementary Fig. S9B), the latter consistent with dysregulation of RB family/E2F regulation.

To assess the potential relevance of these mouse model findings with human genomic data, we compared protein-coding genes with known human orthologs (17, 598 genes) in each group (T-FT, T-O, N-FT, N-O), and selected genes that were uniquely expressed in each to generate group signatures (Supplementary Table S2). Comparison of the mean of scaled expression of the top 50, 100, 250 or 500 T-FT and T-O genes, respectively, with the other four groups showed the expected segregation of samples (Supplementary Fig. S10). We then applied the mouse T-FT and T-O scores (calculated for either the top 50, 100, 250 or 500 DEGs) to TCGA ovarian cancer data. This analysis revealed that human tumors classified as mesenchymal (mean enrichment score=0.27, 0.26 and 0.25 with gene cutoff at Top 100, 250 and 500, respectively. P<0.001 in all cutoffs when compared to other subtypes), and to a lesser extent, as immunoreactive, were more similar to the mouse T-FT, than to the mouse T-O, signature. By contrast, a larger proportion of proliferative type human tumors more closely resembled the mouse T-O signature (mean enrichment score=0.13, 0.16 and 0.15, respectively. P<0.05 in all cutoffs when compared to other subtypes), whereas differentiated tumors segregated fairly equally into those more similar to FTE-derived (mean enrichment score=-0.05, −0.04 and −0.07 with gene cutoff at Top100, 250 and 500, respectively) vs. OSE-derived mouse tumors (mean enrichment score=-0.1, −0.13 and −0.12 with gene cutoff at Top 100, 250 and 500, respectively), respectively (Fig. 6E and Supplementary Table S3). The robustness of the outcome independent of the number of T-FT and T-O genes chosen gives us confidence in the reliability of this analysis. Furthermore, similar results were obtained by analyzing a distinct set of tumor transcriptomes [(22), data not shown)].

### The cell-of-origin can affect response to chemotherapeutic agents

To ask if the cell-of-origin of HGSOC might affect therapeutic response, we tested the response of FTE- and OSE-derived tumorigenic organoids to commonly used ovarian cancer drugs. PTPT (FTE) and LPT (OSE) organoids at day 4 of culture (> 3 passages post-Dox and 4-OHT trea™ent) were released from Matrigel, ∼ 150 organoids were re-seeded in each well of 96-well plates pre-coated with Matrigel, and morphology and viability were assessed 5 days later. Tumorigenic FTE and OSE organoids responded similarly to niriparib, olaparib and gemcitabine (Fig. 7A-C). By contrast, FTE-derived (PTPT) organoids were significantly more sensitive that OSE (LPT) organoids to carboplatin (P<0.05 at 10 µM) and, even more prominently, to paclitaxel (P<0.001 at 5 µM, 10 µM and 25 µM) trea™ent (Fig. 7D and E). These data indicate that the cell-of-origin could influence response to at least some types of chemotherapy.

**Figure 7.**
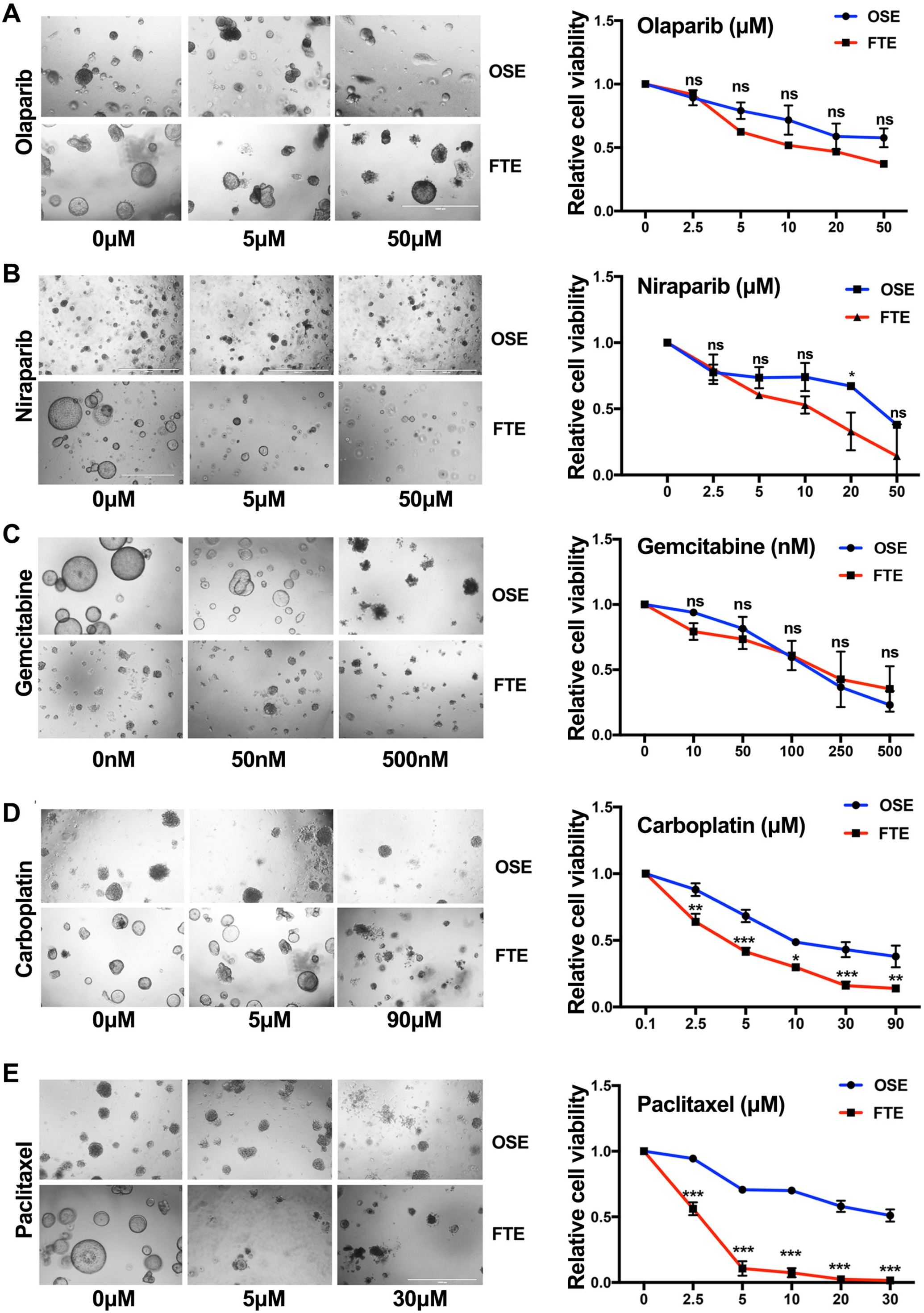
Differential response of FTE- and OSE-derived tumor organoids to chemotherapy. Representative pictures and dose-response curves for LPT (OSE) and PTPT (FTE) organoids, treated with: **A**, Olaparib (0-50 µM), **B**, Niraparib (0-50 µM), **C**, Gemcitabine (0-500 nM), **D**, Carboplatin (0-90 µM) or **E**, Paclitaxel (0-30 µM) on. Cell viability was calculated relative to 0.01% DMSO-treated control cells, measured at 5 days of trea™ent. Each time point represents the means ± SEM of 3 independent experiments, each in triplicate. *p<0.05, ***p < 0.001.

## DISCUSSION

HGSOC is usually diagnosed late, as bulky multifocal disease that has metastasized widely to the peritoneal cavity, a presentation that makes identifying the cell-of-origin difficult (54). Although precursor lesions on the FT fimbriae have been reported in up to 60% of HGSOC cases (13-15), some studies are more consistent with an ovarian or peritoneal origin (5-7). Previous mouse modeling studies showed clearly that FTE can act as the cell-of-origin of HGSOC, but a similar role for OSE has not been excluded. Here, by introducing the same genetic abnormalities into FTE or OSE in GEMMs and organoid models, we have found that HGSOC can originate from either cell type (Supplementary Fig. S11). Importantly, FTE- and OSE-derived tumors differ in their metastatic behavior, transcriptomes and chemosensitivity.

Similar to a previous report, in which combined deletion of *Tp53, Brca1/2, and Pten* was driven by *Pax8rtTA;TetOCre* (37), PTPT mice rapidly developed STIC-like lesions and early (<2 month) metastasis to the ovarian surface. Lineage tracing confirmed that *Pax8 rtTA* is expressed in FTE secretory cells, but importantly, consistent with a recent study (43), we find that *Pax8-*ciliated cells also derive from *Pax8+* progenitors (Supplementary Fig. 2). Thus, while it is clear that HGSOC can arise from mouse FTE, we cannot be certain that FTE *secretory* cells (as opposed to *Pax8-*derived, but *Pax8-* ciliated cells or a specialized *Pax8+* progenitor) are the actual/unique cell-of-origin in the FT. Others have found that the *Ovgp1* promoter also can drive HGSOC (38,39). This *Cre* line purportedly is secretory cell-specific (55), but lineage tracing studies have not been reported. In any event, there should now be little debate that a majority of HGSOC originates from FTE.

Unlike earlier studies using *Pax8rtTA;TetOCre* to drive combined *Brca1/2;Pten;Tp53 deletion*, which gave rise to STIC, invasive HGSOC, and eventually, peritoneal carcinomatosis, PTPT mice died prematurely from massive thymic hyperplasia. Subsequent analysis revealed unexpected, leaky expression of the *TetOCre* promoter in thymic epithelial cells. Previous work showed that E2F activation, resulting from RB family inactivation, leads to increased *Foxn1* expression, which drives thymic epithelial cell proliferation (44). Presumably, *Brca1/2;Pten;Tp53* deletion does not cause such hyperproliferation, although conceivably, these defects could affect other thymic epithelial cell properties and, indirectly, immune function, in such mice. Regardless, our results argue for caution in using the *TetOCre* line to study FTE biology and oncogenesis.

Although HGSOC usually arises from FTE, whether OSE can also give rise to HGSOC had been less clear. Studies in which oncogenic defects were induced by Ad-Cre injections into the ovarian bursa were interpreted as indicating an OSE cell-of-origin (56,57), although others have argued that such injections can result in infection of adjacent tissue, including the oviduct or even the uterus (58). Indeed, our lineage tracing experiments show clearly that bursal injection can target FTE as well as OSE. Nevertheless, several lines of evidence prove that mouse OSE can serve as the cell-of-origin for HGSOC, at least HGSOC induced by combined *Tp53* mutation/inactivation and RB family inactivation. First, Ad-Cre-injected *Tp53*^*R172H/fl*^*;T121* mutant mice develop frank ovarian carcinoma even after salpingectomy. Moreover, *Lgr5-Cre*, which we confirm by lineage tracing marks OSE, but not FTE, in adult mice, also can drive HGSOC generation by combined RB family inactivation/*Tp53* mutation/hemizygosity. Furthermore, OSE organoids in which the above genetic defects are induced *in vitro*, give rise to HGSOC-like lesions.

Although metastasis could not be evaluated in PTPT mice, organoid transplantation clearly shows that mouse FTE-derived HGSOC metastasizes widely. By establishing conditions for OSE-derived organoids, we could directly compare tumorigenesis by FTE- and OSE-derived HGSOC. Interestingly, these experiments suggest that the cell-of-origin can influence HGSOC behavior: FTE-derived tumors showed greater ability to disseminate, whereas OSE-derived HGSOC more commonly formed large, solitary lesions that show less frequent (although nonetheless significant) dissemination. Notably, human HGSOC also differs in its pattern of growth and metastasis, with some cases forming large primary tumors and fewer, less diffuse tumor deposits, whereas others show peritoneal carcinomatosis (59). Defined combinations of genetic defects (e.g., as found in TCGA) can be engineered into our FTE and OSE organoids, allowing various aspects of the transformation process (proliferation, polarity, apoptosis, necrosis, invasion) to be deconstructed *in vitro* and tumorgenicity to be assessed i*n vivo*. It will be interesting to determine the extent to which the differential behavior observed here reflects the particular combination of cell-of-origin/oncogenic defects vs. effects of the cell-of-origin *per se*. Our RNAseq data, however, suggest that whereas common pathways and processes are de-regulated by imposing the same oncogenic insults on FTE and OSE, there are marked differences in which specific genes within these pathways/process are affected in the two cell types. Such differences might manifest as differential sensitivities to the two agents used as front line therapy for HGSOC, carboplatin and paclitaxel. In this regard, it is interesting to note that whereas repair genes are differentially expressed in both FTE- and OSE-derived tumors (compared with their cognate normal tissues), the extent of altered expression is greater in the latter. By contrast, signaling from p53-dependent pathways (presumably by alternative mechanisms, given the combined TP53 mutation/deletion in both types of tumors) is elevated to a greater extent in FTE-derived, than in OSE-derived tumors. Either or both of these differences might help explain the observed cell-of-origin-dependent differences in chemosensitivities.

Several lines of evidence suggest that our results are relevant for human HGSOC. First, a recent case report documented the development of stage IV HGSOC three years post-salpingectomy without evident STIC in the excised FT (60). Furthermore, recent exome sequencing (24), transcriptome (22) and ChiPseq experiments (23) support the notion that HGSOC can arise at sites other than the FTE. Our comparison of the transcriptome of FTE- and OSE-derived HGSOC, respectively, compared with TCGA samples, indicates greater resemblance of some human HGSOCs, particularly proliferative type HGSOC, to OSE-derived tumors. Understanding the contribution of the cell-of-origin to HGSOC generation could be of substantial importance, given that our results indicate that tumorigenic OSE- and FTE-derived organoids bearing the same oncogenic defects have different biologic behavior, including sensitivities to anti-neoplastic drugs.

## METHODS

### Mice

*Rosa26-tdTomato [B6;129S6-Gt(ROSA)26SOR*^*™9(CAG-tdTomato)Hze*^], *[B6;129-Gt(ROSA)26Sor™*^*1sor*^*], Lgr5-121 Cre [B6.129P2-Lgr5*^*™1(cre/ERT2)Cle*^*], Tp53*^*R172H*^ *[B6.129S4(Cg)-Trp53* ^*™2.1Tyj*^*], Trp53*^*flox/flox*^ *[FVB;129-Trp53*^*™1Brn*^ ] and *nu/nu [NU/J]* mice were obtained from the Jackson Laboratory. Conditional *TgK18GT* ^*tg/+*^ BAC transgenic mice (*T121* mice) were described previously (36). *Pax8rtTA* and *TetOcre* strains were described previously (37). *Tp53*^*R172H*^, *Trp53*^*flox/flox*^, *T121* mice were interbred with *Pax8rtTA* and *TetOcre* mice to obtain *Pax8rtTA;TetOcre;Tp53*^*R172H/fl*^ (PTP), *Pax8rtTA;TetOcre;T121* (PTT) and *Pax8rtTA; TetOcre; Tp53* ^*R172H/fl*^*;T121* (PTPT) mice, respectively. *Trp53*^*R172H*^*, Trp53*^*flox/flox*^*, T121* and *Lgr5Cre*^*ERT2*^ were interbred to obtain *Lgr5Cre;Tp53 R172H/fl* (LP), *Lgr5Cre;T121*(LT), *Lgr5Cre; Trp53*^*R172H/fl*^*;T121* (LPT) mice, respectively. *Rosa26-lacz* and *Rosa26-tdTomato* mice were bred to *Pax8rtTA* and *TetOcre* strains to obtain *Pax8rtTA;TetOcre;Rosa26-LacZ* and *Pax8rtTA;TetOcre;Rosa26-tdTomato* mice, respectively. When indicated, mice were euthanized by CO_2_ inhalation and FT and/or ovaries were harvested for histology and organoid culture. All animal experiments were approved by, and conducted in accordance with the procedures of, the IACUC at New York University School of Medicine (protocol no.170602).

### Animal experiments

For lineage tracing of *Pax8+* FTE cells, *Pax8rtTA;TetOcre;Rosa26-LacZ* female mice (6-8 week old) were induced with 2mg/ml doxycycline in their drinking water for 2 days. Mice were sacrificed, and their reproductive systems were collected, at 2 or 60 days after Dox administration. *Lgr5-Cre;Rosa26-tdTomato* mice (6-8 weeks old) were injected intraperitoneally with a single dose of 4-OHT in sesame oil (10 mg/ml) at a concentration of 2 mg (in 200 µl), and then sacrificed at 48h or 4 months after induction. Superovulation was performed by injection with 5 IU pregnant mare serum gonadotropin (PMSG, Sigma) and 5IU of human chorionic gonadotropin (hCG, Sigma), spaced 48 h apart. For *Cre* induction in PTP, PTT and PTPT mice, Dox (2mg/ml) was added for 2 weeks to the drinking water of 6 week-old females, and relevant tissues were collected for H&E and IHC after 1 month. For *Cre* induction in LP, LT and LPT mice, 6 week-old females were injected with 4-OHT, and mice were sacrificed at indicated time.

For ovarian bursa injection, recombinant adenovirus Ad5-CMV-Cre (Ad-Cre) was purchased from the Vector Development Lab at Baylor College of Medicine. Superovulation was performed in 6-8 week old female *Tp53*^*R172H/fl*^*;T121* mice, as described (35). Approximately 1.5 days later, a single injection of 10 µl Ad-Cre (10^11^-10^12^ infectious particles/ml) was delivered into one surgically exposed ovarian bursa, with the contralateral ovary serving as a control. *Rosa26-LacZ* female mice were used to test *Cre* expression and were euthanized 2 weeks post-injection. Salpingectomies on *Tp53*^*R172H/fl*^*;T121* females were performed 3 days after Ad-Cre injection, and mice were euthanized 3 months afterwards. Animals were routinely monitored for signs of distress, poor body condition, and tumor burden, and were euthanized according to veterinary recommendations. For survival experiments, the mice were monitored until their death or upon veterinary recommendation.

Organoids were amplified in 6-well plates coated with 200 µl of Matrigel/well and supplemented with 2 ml organoid growth medium. Orthotopic injections were performed on 8 week-old female mice. For surgeries, mice were anesthetized by intraperitoneal injection of 0.2 ml xylazine hydrochloride, shaved, and cleaned with betadine. A midline dorsal incision was made, followed by an incision into the peritoneal cavity above the fat pad of the right ovary. The ovary was externalized through the incision and 0.5 × 10^6^ cells/matrix mixture (1:1, 15-20 µl) was injected by inserting a 27G needle into the fat pad. Only one ovary in each animal was injected, with the contralateral ovary serving as a control. The injected tissue was returned to the peritoneal cavity, the inner incision was sutured, and the outer incision was sealed with wound clips. For mammary fat pad injections, cells were resuspended with Matrigel (1:1, around 15 µl) and injected into the mammary fat pad just inferior to the nipple of female mice (6-8 weeks) with 28G needle (BD insulin syringe). Mice that developed tumors were euthanized at the indicated times, when tumors ulcerated or reached a maximum diameter of 20 mm, or when mice showed any sign of discomfort due to tumor burden.

### Organoid Cultures and Assays

For FTE organoids, FT fimbriae from wild type, PTP, PTT and PTPT were dissected under a microscope, minced and digested with Collagenase type I and dispase 0.012% (w/v) (STEMCELL Technologies) at 37°C for 1 hour, followed by incubation in TrypLE(tm) Express Enzyme (Thermo Fisher Scientific) for 10 min at 37°C, and inactivation with FBS (Gibco) 1% in DMEM media (Gibco). Dispersed FTE cells were passed through a strainer (70 μm), mixed with Matrigel (BD Bioscience), seeded and cultured as described(61). After Matrigel solidified (10 min at 37 °C incubator), culture medium was added. The medium was based on Ad+++ medium (AdDMEM/F12 (Invitrogen) with HEPES (Thermo Fisher Scientific), penicillin/streptomycin (Life Technologies), Glutamax (Life Technologies)), supplemented with B27 (Invitrogen), N2 supplement (Thermo Fisher Scientific), 1.25 mM N-acetylcysteine (Sigma), 50 ng/ml EGF (Thermo Fisher Scientific), 500 ng/ml RSPO1 (Peprotech) or R-spondin-1-conditioned medium (25%, v/v), and 100 ng/ml Noggin (Peprotech). For the first 3 days after thawing, culture medium was supplemented with 10 mM Y-27632 (Sigma-Aldrich). For OSE organoid culture, fat and adjacent fallopian tubes were carefully removed under a microscope, and the ovaries were digested with 0.25% trypsin/EDTA (Invitrogen) and inactivated with DMEM media containing 1% FBS (Gibco) at 37°C for 30 min. Supernatants, containing cells stripped from the OSE, were seeded in Matrigel, and cultured in Ad+++ medium, supplemented with B27, N-acetylcysteine (1 mM, Sigma), WNT3a-conditioned medium (50% v/v), R-spondin-1-conditioned medium (10% v/v), Noggin (100 ng/ml, Peprotech), EGF (0.0125 µg/ml), nicotinamide (10 mM, Sigma), A83-01 (0.5 µM, Tocris Bioscience), hydrocortisone (0.5 µg/ml, Sigma), and β-estradiol (100 nM, Sigma). Passaging and freezing were carried out as described above for FTE organoids. WNT3a- and R-spondin-1-conditioned media were prepared as described (62).

Organoid medium was changed every 2-3 days, and they were passaged approximately 1:4 (10,000 cells/well) every 6-8 days. For passage, growth medium was removed, and the Matrigel was resuspended in cold Cultrex^®^ Organoid Harvesting Solution and transferred to a 15-ml Falcon tube, which was placed on ice for 15 minutes. Organoids were recovered by centrifugation at 1,000 g for 5 minutes and resuspended in 500 µl TrypLE Express Enzyme (Gibco) for 10 minutes at 37°C. Cells were then seeded as indicated for each experiment. For freezing, cells were resuspended in organoid medium with 10% DMSO and 10% FBS, cooled, and stored in liquid nitrogen.

To activate *Cre in vitro*, 0.5 µl of Dox (1 mg/ml) were added to 500µl of freshly isolated PTP, PTT and PTPT cells, which were plated as described above. Similarly, 1 µg/ml 4-OHT was used to activate CRE in LP, LT and LPT organoids. All data were generated at least 3 passages after induction. Organoid size was quantified as the surface area of horizontal cross sections. If all organoids in a well could not be measured, several random, non-overlapping pictures were acquired from each well using an Invitrogen(tm) EVOS(tm) FL Digital Inverted Fluorescence Microscope, and analyzed by using ImageJ software. Organoid perimeters for area measurements were defined manually and by automated determination using the “Analyze Particle” function of ImageJ, with investigator verification of the automated determinations. Organoids touching the edges of images were excluded from counting.

To compare organoid-forming efficiency of different genotypes, 5,000 cells were seeded into a 24-well plate, total organoid number was counted under a light microscope after 5–7 days in culture, and the percentage of organoids formed relative to organoids formed by wild type cells was calculated.

For *in vitro* growth curves, organoids were incubated in TrypLE Express (Gibco) for 15 min at 37°C, followed by an additional 5 min digestion in Dispase I. Isolated cells were passed through a strainer, seeded at 2 ×10^4^ cells/well in a 24 well plate and placed in culture medium. At the indicated times, cells were recovered as described above, and viable counts were obtained by trypan blue exclusion.

Drugs were tested in organoids based on a previously described protocol (62). Briefly, organoids in culture at day 4 were released from Matrigel and diluted to 50 organoids/µl in growth medium lacking N-acetylcysteine and Y-27632. Clear bottom 96-well plates were coated with 20 µl Matrigel before the addition of 30 µl of organoid suspension. The indicated concentrations of paclitaxel (Selleck Chem), carboplatin (Sigma), olaparib (Selleck Chem), niraparib (Selleck Chem), or DMSO (control) were added in triplicate. On day 5 of trea™ent, media were removed and the Matrigel drops were suspended in 40 µl CellTiter-Glo 3D (Promega) and 80 µl advanced AdDMEM/F12, incubated for 30 min at room temperature before luminescence was measured in FlexStation^®^ 3 Multi-Mode Microplate Reader. Results were normalized to DMSO controls.

Invasion was assessed by using chambers with 8 µm pore size polycarbonate membrane (Transwell) inserts (Costar). Matrigel (30 µl) was added to the chambers and allowed to solifiy at 37C for 10 minutes. Wild type, PTP, PTT, or PTPT cells (2 × 10^4^/50µl Ad+++ medium/well) were seeded into the Matrigel-coated top chamber, and allowed to attach for 12 hours, followed by the addition of 500 μl of culture medium to each well. After an additional 96 hours of incubation, the upper surface of the Transwell membrane was scrubbed carefully several times with a cotton swab soaked in PBS to remove non-invaded cells. The lower membrane was then rinsed with PBS carefully several times, and cells that had invaded were visualized by staining with crystal violet and counted under a microscope. Invasion was calculated as the average number of cells per 10x field, determined by Image J software.

### FACS

Ovaries from Lgr5-Cre (*Lgr5–EGFP–ires–Cre*^*ERT2*^) females (6-8 weeks) were digested as described above and recovered OSE cells were passed through a strainer (40 μm) to obtain single-cell suspensions. OSE cells were pelleted by centrifugation at 1,000 g for 5 min and resuspended in PBS containing 2% FBS, Rock inhibitor (Y-27632, 10 µM, STEMCELL Technologies Inc.), DAPI (1µg/ml), FACS was performed immediate by using a MoFlo^™^ XDP, and sorted GFP^hi^ and GFP^neg^ cells were seeded at 5000/well. Organoids were counted 6 days later, and organoid forming efficiency was calculated as number of the organoids/number of cells seeded, pooled from three biological replicates.

### Histology and Immunostaining

Tissues were fixed in 4% paraformaldehyde (PFA) in PBS at 4°C for 4 hours. Organoids were fixed in 4% PFA for 15 minutes and placed in Histogel (Thermo Fisher Scientific) before tissue processing and embedding. Formalin-fixed paraffin-embedded (FFPE) tissue sections (5 µm) were de-paraffinized, rehydrated, and then stained with hematoxylin and eosin (H&E) or subjected to IHC. For antigen retrieval, slides were autoclaved in 0.01 M citrate buffer (pH 6.0). Endogenous peroxidase activity was quenched in 3% H_2_O_2_ in methanol for 15 min and sections were blocked with 0.5% BSA-PBS for 1h. Primary antibodies were added overnight at 4°C, then washed with PBS 3×10 min, incubated with HRP-labeled secondary antibodies, and after washing with PBS 3x for 10 minutes each, signals were visualized by using the HRP Polymer Detection Kit and DAB peroxidase (HRP) substrate (34002, Life Technologies). Primary antibodies included: Ki67 1:200 (ab15580, Abcam), γ-H2AX 1:500 (05-636, Thermo Fisher Scientific), CK7 1:200 (ab181598, Abcam), Stathmin-1 1:200 (3352S, cell signaling), P16 1:200 (sc-1661, Santa Cruz), PAX8 1:200 (10336-1-AP, Proteintech), p53 1:800 (P53-CM5P-L, Leica). Secondary antibodies included: goat anti-chicken IgY-HRP 1:200 (sc-2428, Santa Cruz), goat anti-rabbit IgG-HRP 1:200 (sc-2030, Santa Cruz).

Immunofluorescence was performed in frozen tissue sections (5 µm) or in whole organoids released by gently dissolving the Matrigel in ice-cold PBS. Following fixation as above, cells were permeabilized in PBS containing 0.5% Triton X-100 and blocked in PBS containing 1% BSA, 3% normal goat serum and 0.2% Triton X-100. Primary antibodies were incubated at 4°C overnight, and sections were washed in PBS 3x for 10 minutes each, followed by incubation with DAPI (2 µg/ml) and Alexa 488-, 555- or 647-conjugated anti-chicken, anti-rabbit or anti-mouse antibodies, as indicated. After washing, samples were mounted with ProLong Gold antifade reagent (Life Technology). Primary antibodies were: GFP 1:300 (ab13970, Abcam), WT1 1:200 (ab15249, Abcam), E-cadherin 1:200 (ab15148, Abcam). Secondary antibodies included: goat anti-mouse IgG, Alexa Fluor^®^ 647 conjugate 1:200 (A28181, Thermo Fisher Scientific), goat anti-rabbit IgG, Alexa Fluor^®^ 555 conjugate 1:200 (A27039, Thermo Fisher Scientific), and goat anti-chicken IgY H&L (Alexa Fluor^®^ 488) 1:200 (ab150169, Abcam).

### Laser capture microdissection and RNA extraction

Tissues, including normal ovary and fallopian tube (fimbriae), ovarian tumors from LPT females, and STICs from PTPT mice, were embedded in FSC 22 Clear Frozen Section Compound and immediately frozen with liquid N_2_. The resultant blocks were cut at 5-8 µm and mounted on a PEN membrane frame (Leica), followed by air-drying the slides for 30 minutes at room temperature. Laser capture was performed with a Leica LMD6000 laser microdissection system, and excised pieces were harvested into 200 µl RNase-free tubes, which were carefully recovered from the microscope, centrifuged, and placed on ice. RNA was extracted by using the miRNeasy mini Kit (Qiagen), following the manufacturer’s instruction.

### RNA-Sequencing and Analysis

Libraries were prepared using the Illumina TruSeq Stranded Total RNA Sample Preparation Kit. and sequenced on an Illumina HiSeq 4000 sequencer using 150bp paired-end reads by the Perlmutter Cancer Center Genome Technology Center shared resource (GTC). Sequencing results were demultiplexed and converted to FASTQ format using Illumina bcl2fastq software. The average number of read pairs was 60.3M per sample. Data were processed by the Perlmutter Cancer Center Applied Bioinformatics Laboratory shared resource (ABL). The reads were adapter and quality trimmed with Trimmomatic [http://www.ncbi.nlm.nih.gov/pubmed/24695404] and then aligned to the mouse genome (build mm10/GRCm38) using the splice-aware STAR aligner [http://www.ncbi.nlm.nih.gov/pubmed/23104886]. The featureCounts program [https://www.ncbi.nlm.nih.gov/pubmed/24227677] was utilized to generate counts for each gene based on how many aligned reads overlap its exons. These counts were then normalized and used to test for differential expression using negative binomial generalized linear models implemented by the DESeq2 R package [http://www.ncbi.nlm.nih.gov/pubmed/25516281]. Statistical analysis and visualization of gene sets were performed using the clusterProfiler R package [https://www.ncbi.nlm.nih.gov/pubmed/22455463].

FT/O-specific gene signatures were determined by calculating differentially upregulated genes pair-wise between all four examined groups. For each group, they were ranked by the highest p-value across all three comparisons for each gene. The genes were subset to protein coding and with available human ortholog predictions in any of the 14 databases indexed by the HUGO Gene Nomenclature Committee Comparison of Orthology Predictions resource. To measure the FT/O enrichment in individual samples, the normalized counts were scaled and centered, and the mean was calculated for all genes comprising the signature. TCGA Ovarian Cancer batch effects normalized mRNA data and molecular subtypes were retrieved from the UCSC Xena Pan-Cancer Atlas hub.

### Quantification and statistical analysis

Unless otherwise specified, data are presented as mean ± SEM. Survival rates were analyzed by log-rank test, using GraphPad Prism software. P values were determined by two-tailed Student’s t-test, unless otherwise specified, with P< 0.05 considered statistically significant.

## Acknowledgments

We thank Terry Van Dyke for sharing the T121 mice. We would also like to acknowledge technical support for this work provided by the PCC Experimental Pathology, Microscopy, GTC and ABL shared resources (P30CA016087). We also thank Drs. Jiyuan Hu (Division of Biostatistics), Chan Wang (Division of Biostatistics), Kwan Ho Tang (PCC) and Victor Ho (Princess Margaret Cancer Center) for advice and discussion. Initial work on this project was supported by grant-MOP-191992 from the Canadian Institutes for Health Research to B.G.N. D.A.L is supported by the Depar™ent of Defense Ovarian Cancer Research Program (W81XWH-15-1-0429), and S. Z. was supported by a post-doctoral fellowship from the Ovarian Cancer Research Fund Alliance.

## SUPLLMENTARY FIGURE LEGENDS

**Supplementary Figure 1:**
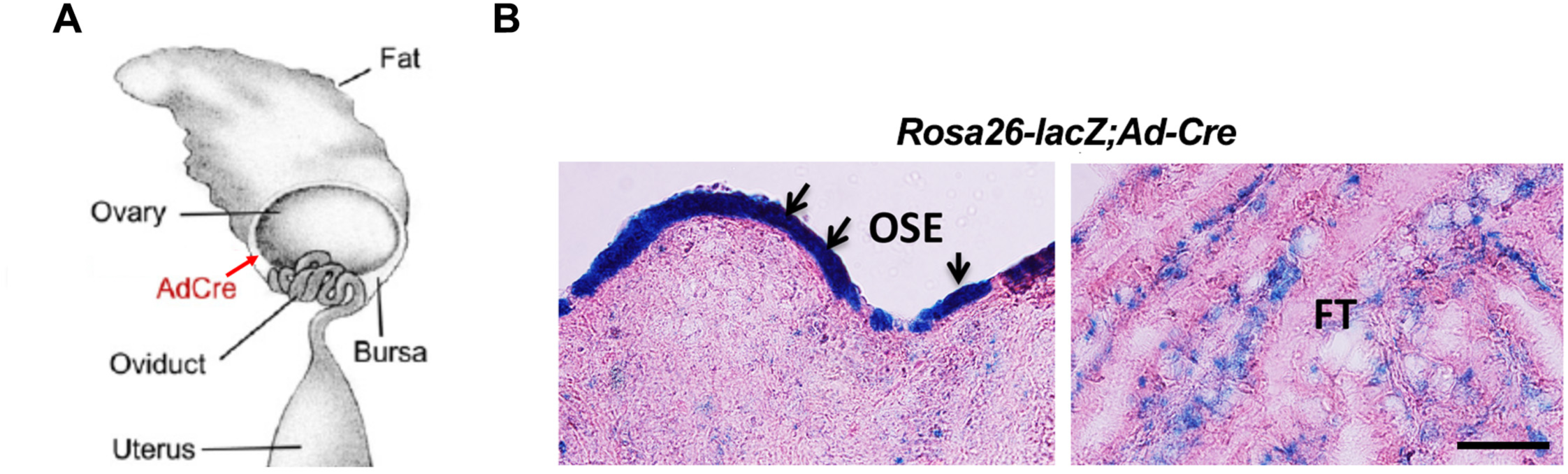
Ad-Cre injected into the ovarian bursa targets OSE *and* FTE. **A,** Schematic showing AdCre injection into ovarian bursa. **B**, X-gal staining shows β-galactosidase expression in OSE (left) and FTE (right) following injection.

**Supplementary Figure 2:**
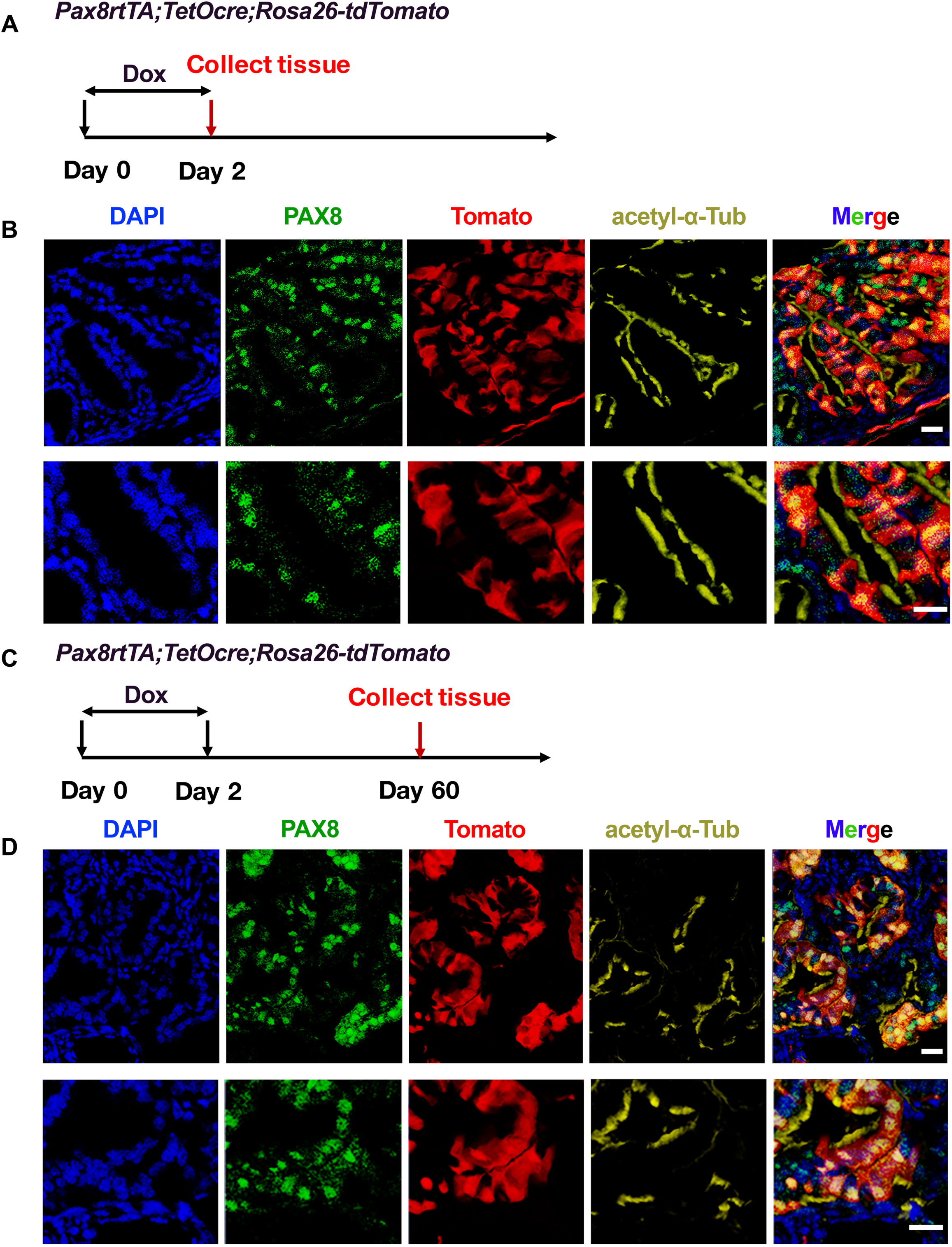
Lineage tracing of *Pax8* cells in the mouse fallopian tube. **A,** Scheme depicting induction of adult *Pax8-rtTA;tetO-Cre;Rosa26-tdTomato* mice by trea™ent with Dox for 2 days, followed by sacrifice and tissue collection. **B,** Immunofluorescence staining for acetylated-α-tubulin (yellow), PAX8 (green) and DAPI (blue) in mouse oviductal epithelium after 2-day Dox trea™ent. **C,** Scheme showing chase experiment. **D**, Immunofluorescence staining for acetylated-α-tubulin (yellow) and PAX8 (green) in oviductal epithelium of *Pax8rtTA;TetOcre;Rosa26-tdTomato* mice, 60 days after Dox induction. The lower panels in (**B**) and (**D**) represent a higher magnification of the corresponding upper panel. DAPI-stained nuclei are shown in blue. Tomato (red), acetylated-α-tubulin (yellow), PAX8 (green) and DAPI (blue) staining, with the overlap (Merge) shown in the right panels. Scale bars: 20 µm.

**Supplementary Figure 3:**
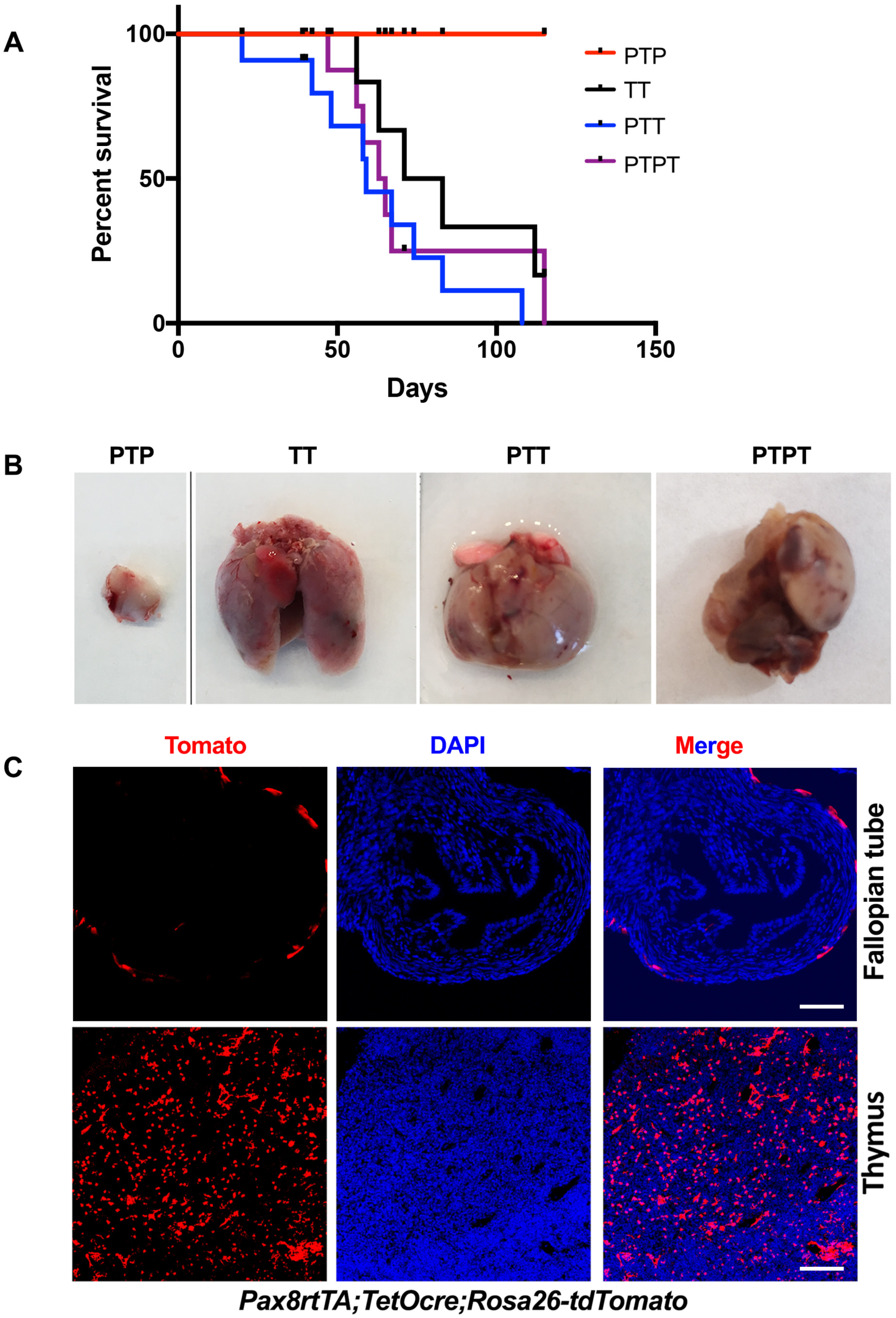
“Leaky” expression of *TetOCre* in thymic epithelium of PTT or PTPT mice results in lethal hyperplasia. **A,** Survival curves of PTP, TT, PTT and PTPT mice. TT: *TetOcre;T121***. B,** Representative morphology of thymi from PTP, TT, PTT and PTPT mice. **c,** Representative Tomato immunofluorescence in thymi from *Pax8rtTA;TetOcre;Rosa26-tdTomato* without Dox trea™ent. Slides were counterstained with DAPI. Scale bars: 10 µm.

**Supplementary Figure 4:**
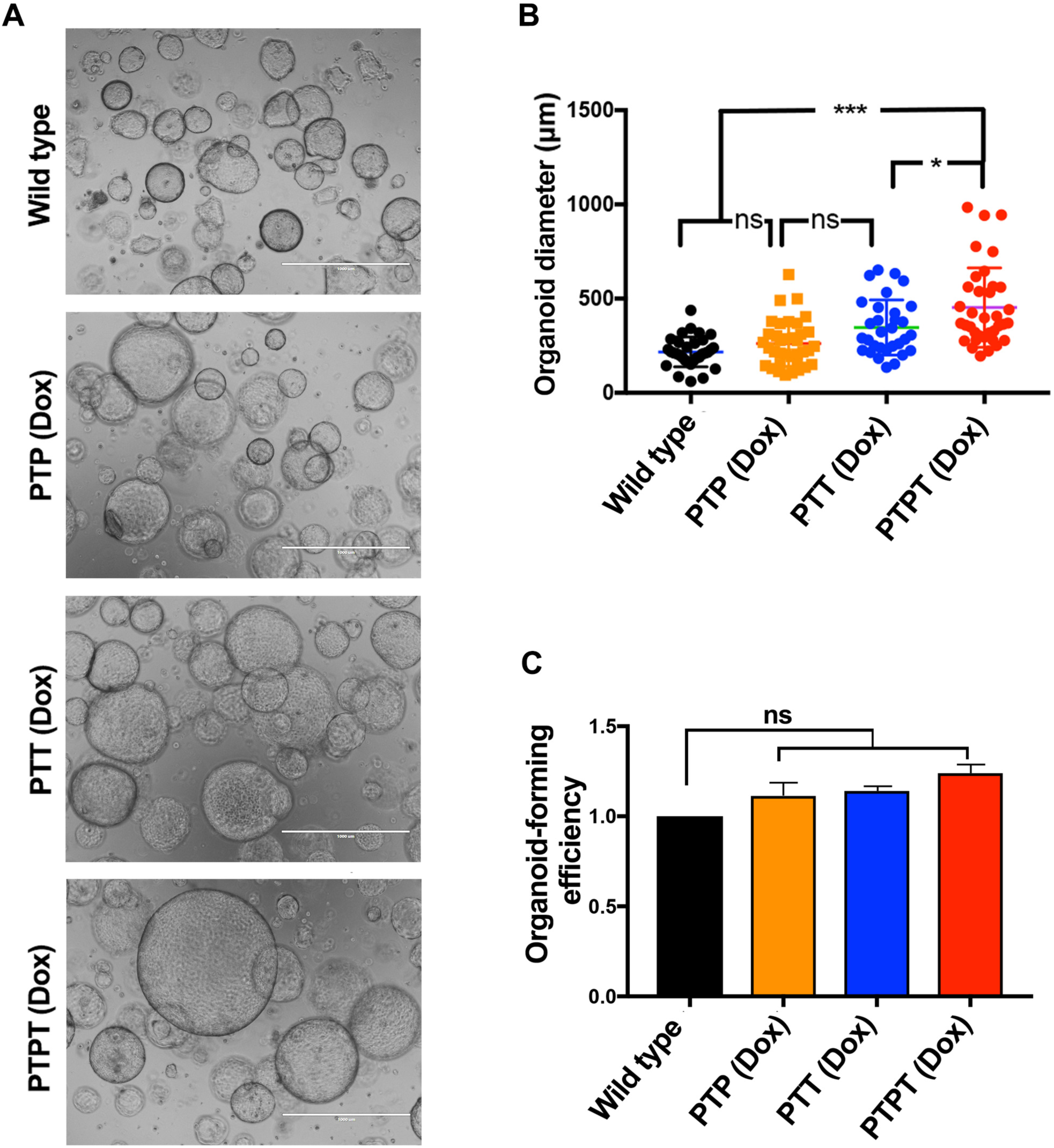
Effect of mouse genotype on FTE organoid size. **A**, Bright field views of FTE organoids from the indicated mice, cultured for 6 days. Each culture was established by seeding 5,000 FTE cells. **B**, Average diameter of organoids from the indicated genotypes, measured on day 6 of culture. Organoids from 3 wells were analyzed for each group. Data represent means ± SEM. *P<0.05; ***P<0.01. **C**, Organoid-forming efficiency for FTE cells from the indicated genotypes, quantified at day 6 of culture, from organoids derived from 3 independent mice of each genotype. Data represent the mean ± SEM. Statistical analysis was performed by Dunnett’s multiple comparison test. ns, not significant.

**Supplementary Figure 5:**
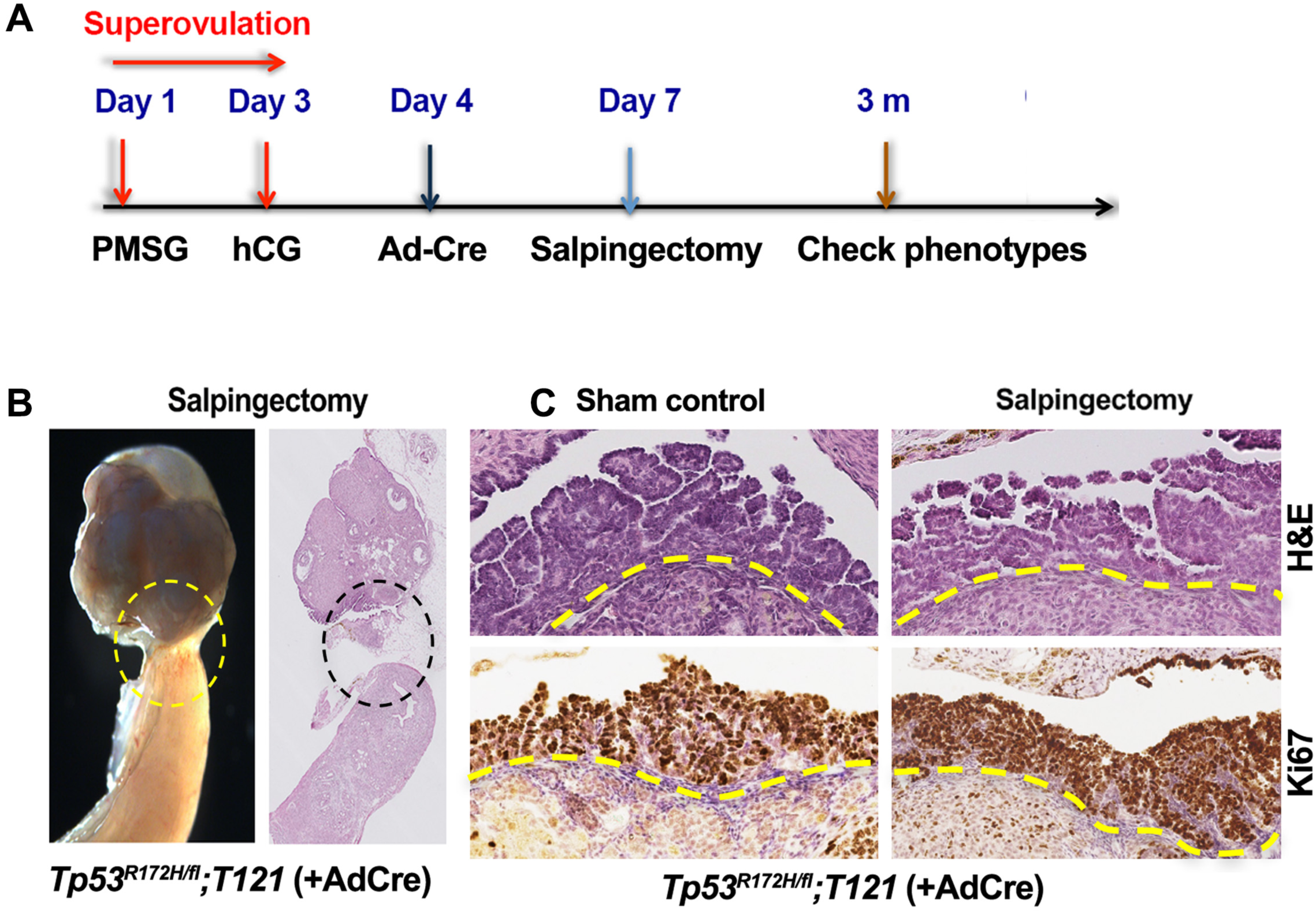
Salpingectomy does not prevent neoplasia in Ad-Cre-induced *Tp53*^*R172H/fl;*^*T121* mice. **A**, Schematic showing experimental strategy. **B**, Micrograph showing gross morphology (Left panel) and H&E-stained section of female genital tract from salpingectomized *Tp53*^*R172H/fl*^*;T121* mice, injected with Ad-Cre (Right panel). Dashed yellow ovals show lack of fallopian tube in the female genital tract. **C,** Representative H&E stain and Ki67 IHC of section from Ad-Cre-injected *Tp53*^*R172H/fl*^*;T121* mice with or without (Sham) salpingectomy, assessed 3 months post injection. The yellow dashed line shows the border between tumor and underlying cells.

**Supplementary Figure 6:**
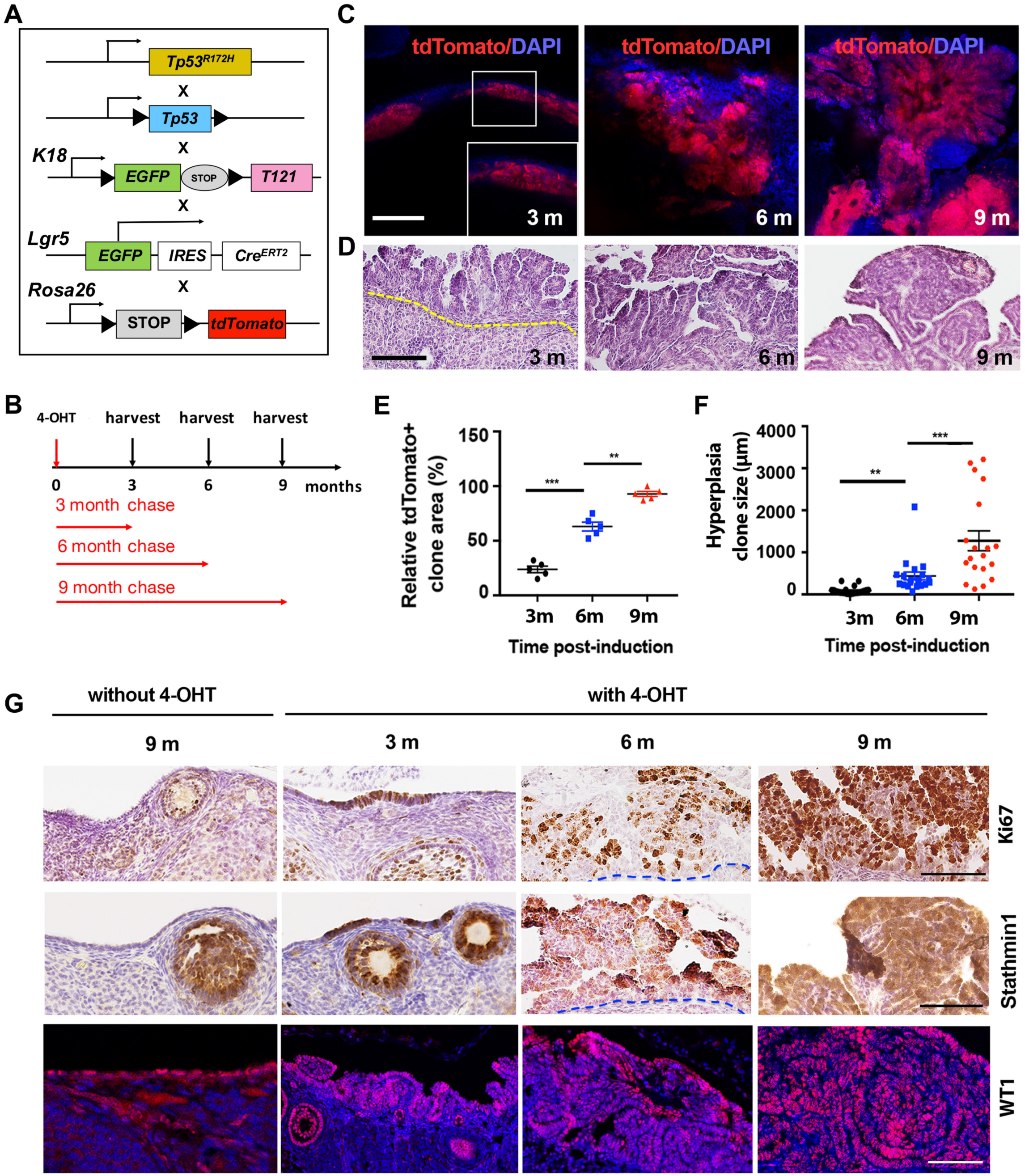
*Lgr5 +* cells can initiate tumors with markers of HGSOC. **A,** Schematic depicting crosses of *Tp53*^*R172H/-*^*;T121;Lgr5Cre* and *Rosa26-tdTomato* mice**. B,** Lineage tracing scheme for *Lgr5Cre;Rosa26-Tdtomato* mice. **C,** Whole mounts of ovaries from *LPT;Rosa26-Tdtomato* mice induced with 4-OHT at 6 weeks, followed by the indicated chase times, showing Tomato (red) and DAPI (blue) fluorescence. In left panel, region within the top white box is magnified below. Note expansion of isolated Tomato+ cells into Tomato+ clones. Scale bar, 50 µm. **D,** Corresponding H&E-stained sections from ovaries in (**C**). The yellow dashed line shows the border between the OSE-derived tumor and underlying cells. **E,** Relative areas of Tomato+ OSE clones at the indicated times after 4-OHT induction (“chase times”). Clone size was determined by measuring the longest ‘length’ of a discrete Tomato+ clone. Each circle represents a distinct Tomato+ clone, with n=3 ovaries analyzed/time point, and a total of 180 clones counted. **F,** Quantification of hyperplastic clone size on the ovarian surface. Clone size was determined by measuring the longest ‘length’ of epithelial protrusions (tumor cells) from the border with stromal cells (yellow dash line in **D**). Twenty (20) cross-sections from 3 ovaries were counted for each time point. Data represent mean ± SEM. *p < 0.05, **p < 0.01, and ***p < 0.001. **G,** Representative IHC (p53, Ki67 and Stathmin-1) and immunofluorescence (WT-1) staining for key HGSOC markers in OSE from LPT mice at the indicated times after 4-OHT induction, or without trea™ent.

**Supplementary Figure 7:**
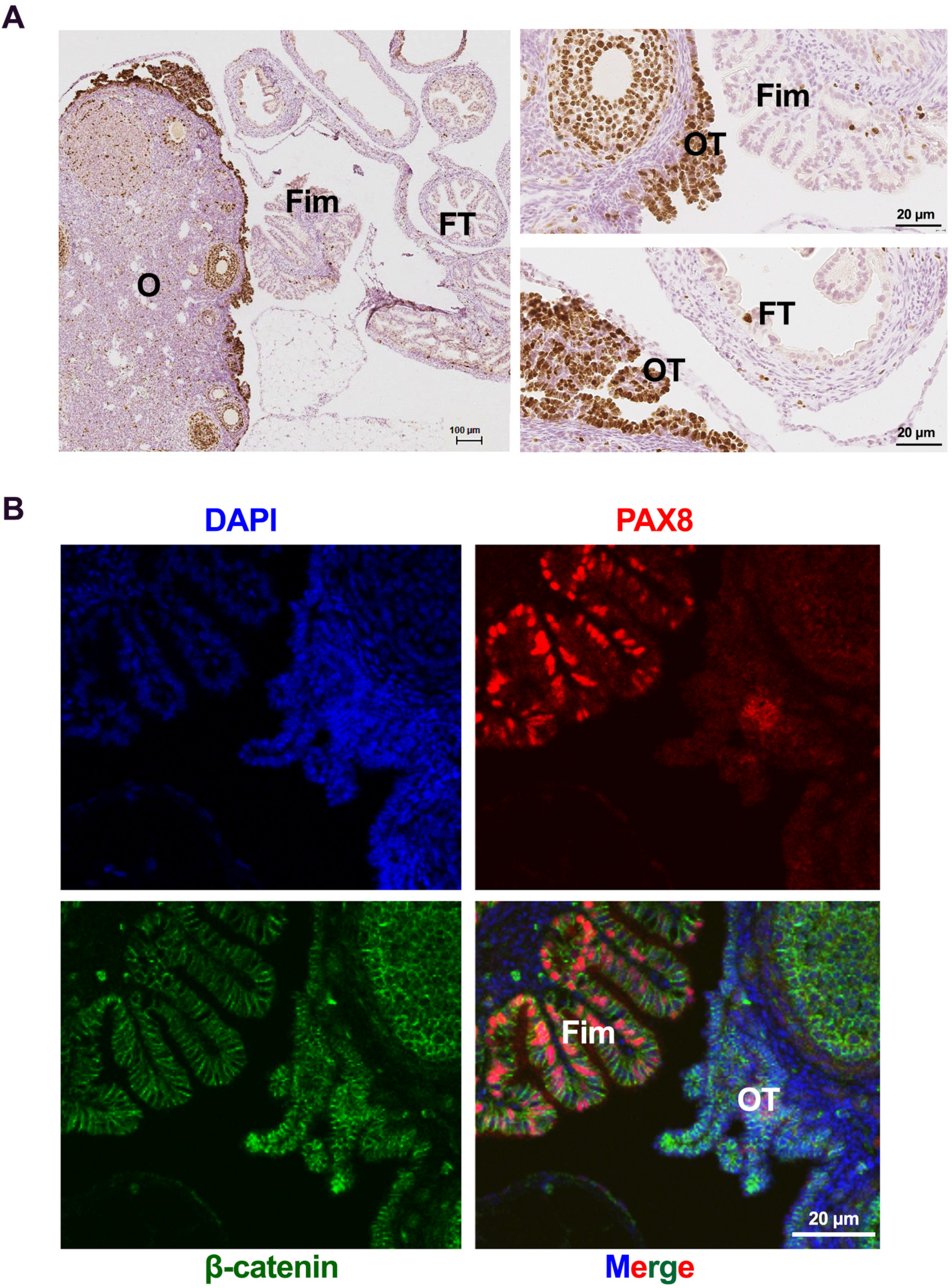
*Lgr5-Cre* only directs tumorigenesis in OSE. **A,** Ki67 IHC staining of OSE and FTE from an LPT mouse. Right panels are higher magnifications (20X) of the left image. **B**, Immunofluorescence staining for PAX8 (red) and β-catenin (green) in ovary and fallopian tube, DAPI (blue) was used as nuclear counterstain. Note that FTE stains strongly for PAX8, while OT is PAX8 negative. OT: OSE-derived tumor. FT: fallopian tube. Fim: fimbria.

**Supplementary Figure 8:**
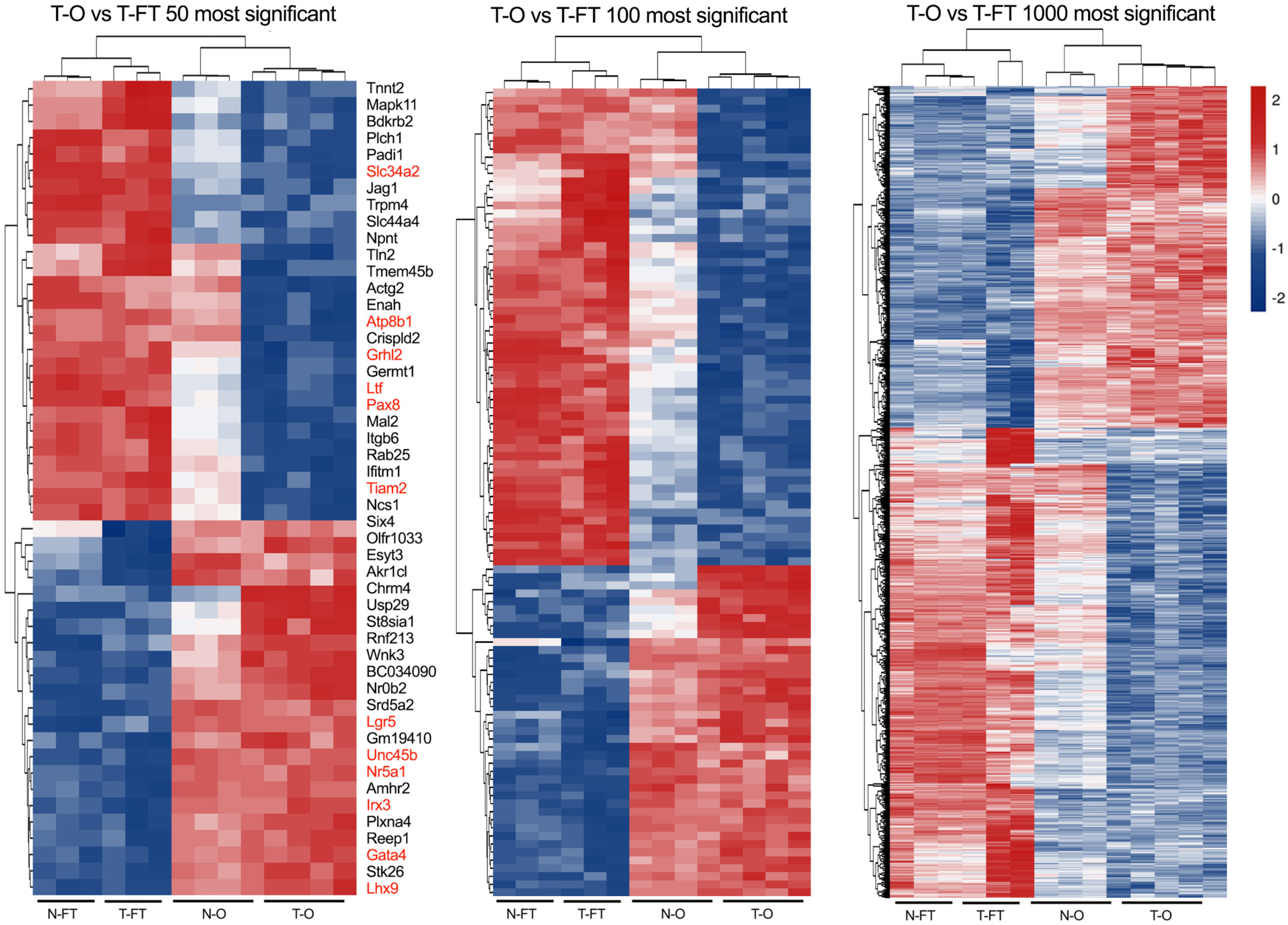
Contribution of lineage-dependent and -independent gene expression to differential gene expression in FTE-vs OSE-derived tumors. Hea™ap of the 50, 100 and 1000 most significant differentially expressed genes in OSE-derived (T-O) and FTE-derived (T-FT) tumors. For reasons of space, gene names are shown only for the top 50 genes. Red colored genes are reported lineage-related genes.

**Supplementary Figure 9:**
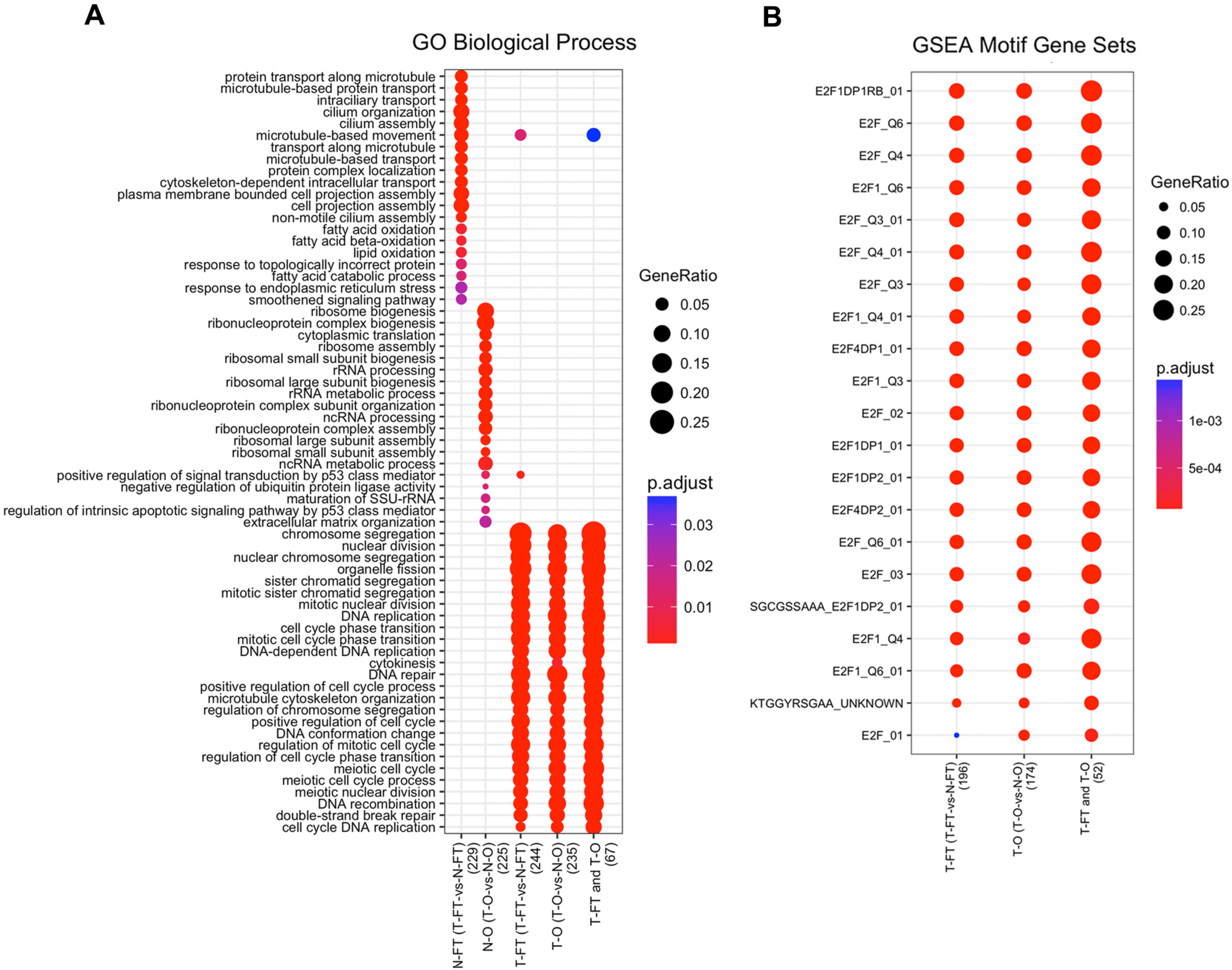
Additional pathway analyses comparing normal FTE and OSE and the tumors derived from each. **A,** GO categories analysis **B,** Gene set enrichment analysis (GSEA). The size of each circle represents the number of DEGs within each category; color coding indicates the significance level.

**Supplementary Figure 10:**
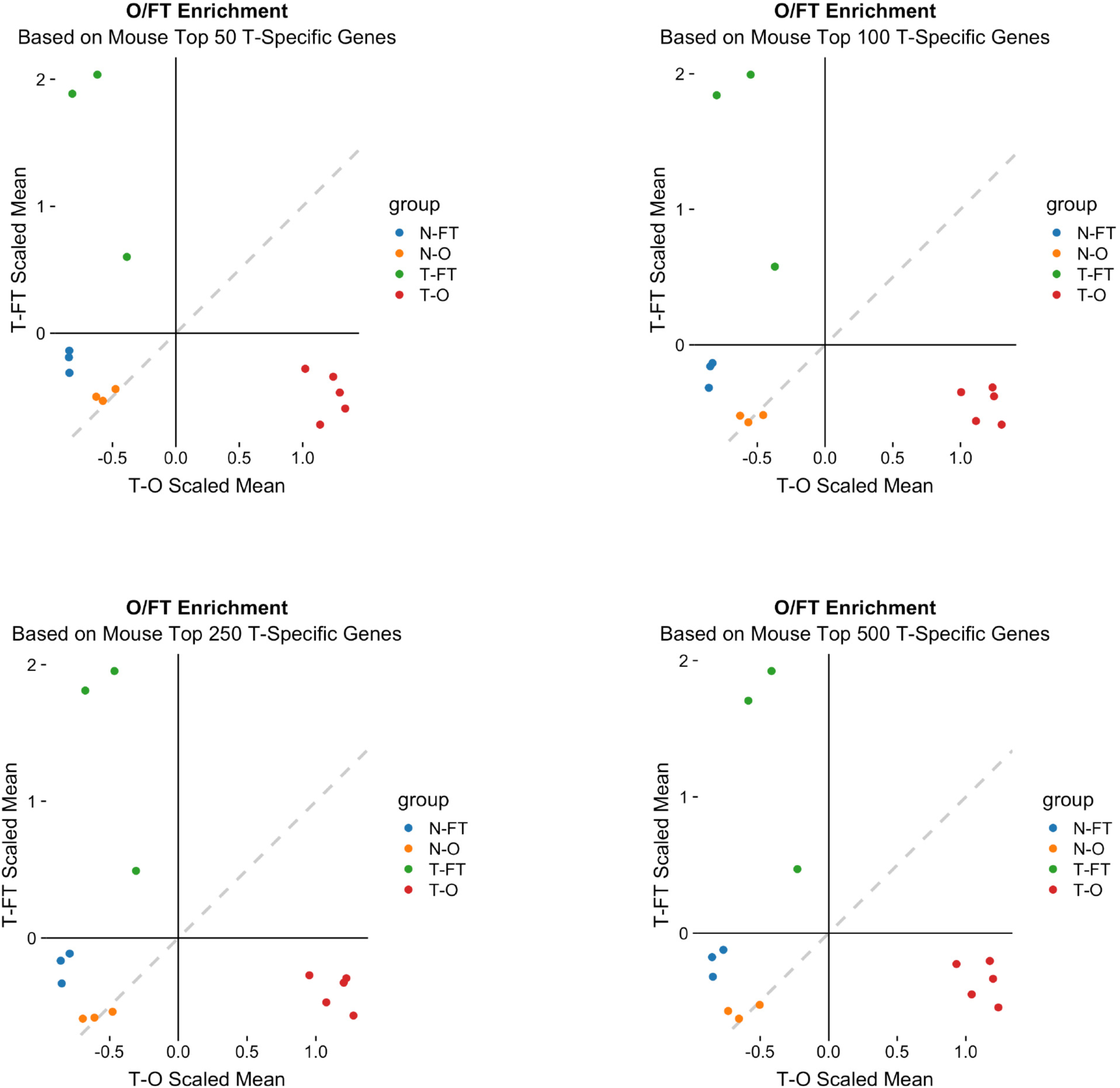
Validation of ovarian and fallopian tube-derived tumor signatures. Signatures were developed based on the top 50, 100, 250, or 500 FTE- or OSE-derived tumor-specific genes, and compared with the mean scaled gene expression from normal FTE (N-FT), normal OSE (N-O), and FTE (T-FT)- or OSE (T-)-derived tumors, as indicated. See Methods for details.

**Supplementary Figure 11:**
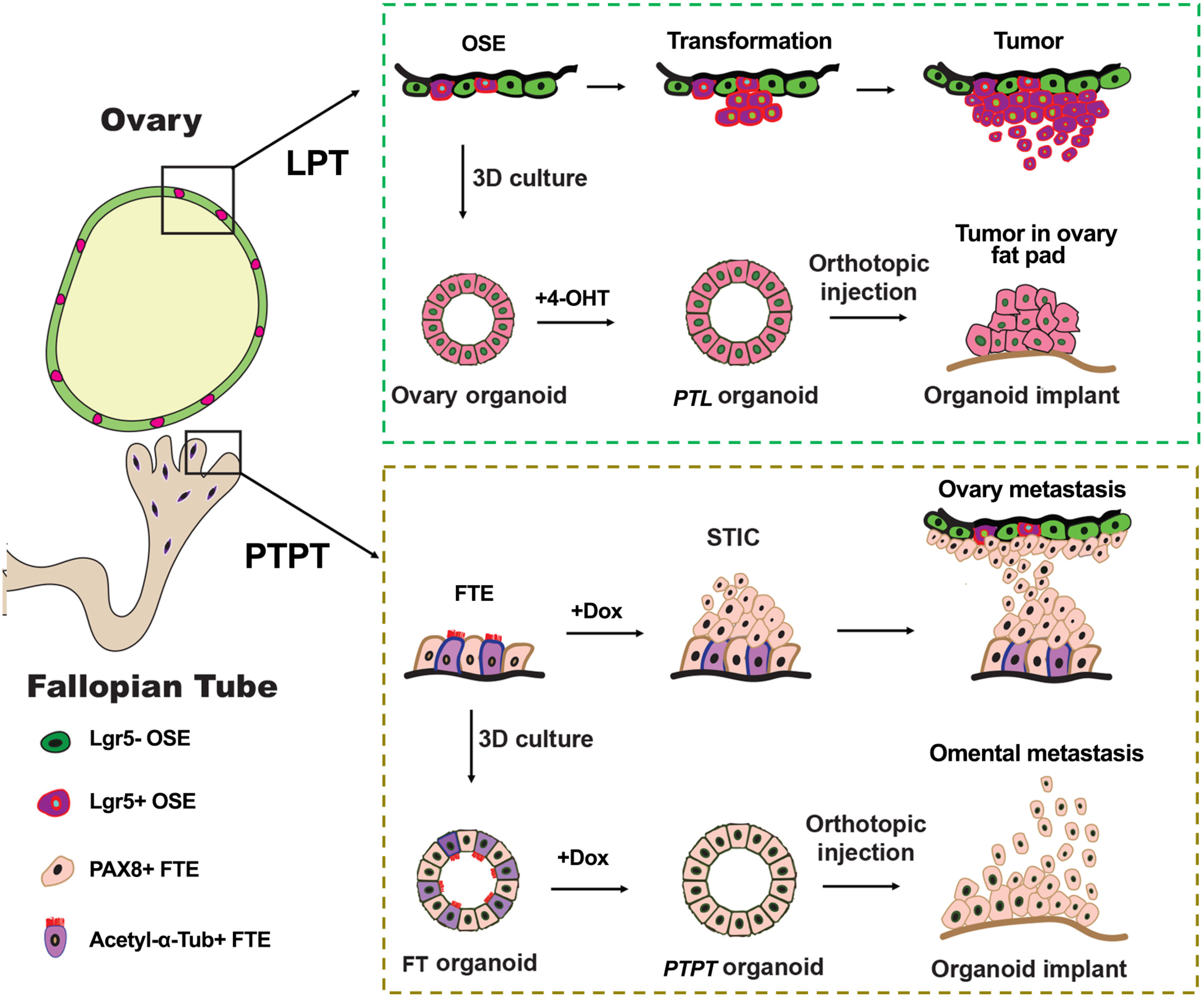
Model showing approaches used to assess cell-of-origin of HGSOC. Upper Panel: Combined RB family inactivation (via T121 expression) and *Tp53* mutation in *Lgr5+* OSE cells, initiated by CRE activation (with 4-OHT), results in patchy areas of OSE transformation that expand, taking over the entire ovarian surface and eventually spreading to the peritoneal cavity. These processes can be reproduced by transplanting cognate OSE organoids. **Lower Panel**: The same genetic events also cause transformation of *Pax8+* FTE, leading to Serous Tubal Intraepithelial Carcinoma (STIC) and ovarian metastasis. These behaviors also can be recapitulated in FTE organoids, which generate widespread abdominal metastasis following orthotropic injection.

**Supplementary Table S1: RNA analyses. A,** DEGs shared between N-O vs N-FT and T-O vs T-FT at Pvalue<0.05. **B,** Top50 DEGs between T-O (OSE-derived tumor) and T-FT (FTE-derived tumor), N-O (Normal OSE) and N-FT (Nomal FTE) at a fold change (FC) >2 (log2FC > 1). **C,** DEGs in diferent comparisons in T-O vs N-O, T-FT vs N-FT and the shared DEGs between T-O vs N-O and T-FT vs N-FT. All at Pvalue<0.05.

**Supplementary Table S2: Top 100, 250 and 500 T-FT and T-O specific mouse genes and the corresponsive HGSOC patient genes from TCGA.**

**Supplementary Table S3: Scores and P values of multiple comparisons among T-FT and T-O enriched HGSOC subtypes.**

